# Adaptive epigenetic regulation of neuronal metabolism by a mitochondrial redox signal

**DOI:** 10.1101/2023.12.10.570533

**Authors:** Marius W. Baeken, Philipp Kötzner, Holger Richly, Christian Behl, Bernd Moosmann, Parvana Hajieva

## Abstract

Different signaling pathways connect the mitochondrion with the transcriptional machinery in the nucleus. Redox events are thought to play a substantial role along this axis, however, many open questions about their specificity, quantitative importance and mode of action remain. Here, we have employed subtoxic doses of the complex I inhibitor MPP^+^ in human neuronal LUHMES cells to characterize the contribution of scavengeable redox signals to mito-nuclear communication. MPP^+^ evoked a broadly targeted transcriptional induction of nuclear-encoded respiratory chain complex (RCC) subunits. Nanomolar doses of phenothiazine (PHT), a mitochondrially active antioxidant, attenuated these transcriptional effects by approximately half, but did not modulate the bioenergetic markers ATP, NAD^+^, NADH, lactate, or glucose. Transcriptional induction by MPP^+^ was accompanied by a loss of nuclear 5-methyl-cytosine and an increase in histone H3K14 acetylation, both of which were entirely prevented by PHT. Inhibitor and PHT reversibility experiments suggested that these alterations were mediated by lowered DNMT3B and SIRT1 levels, respectively. Analysis of MPTP-treated mice recapitulated the PHT-reversible induction of histone acetylation and DNMT3B suppression in vivo. Moreover, PHT completely abrogated the statistical significance of the association of MPP^+^ with the selective induction of mitochondrially imported proteins and RCC subunits. We conclude that the mitochondrion employs a redox signal to announce impending, but not yet acute mitochondrial distress to the nucleus, in order to selectively upregulate mito-metabolic genes via chromatin reorganization. Our results have implications for the interpretation of the observed epigenetic changes in Parkinson’s disease and other neurodegenerative disorders.

## Introduction

NADH dehydrogenase is the first complex of the canonic mitochondrial respiratory chain (Lenaz and Genova, 2009; Brand, 2016). Isolated complex I deficiency presents as energy generation disorder that frequently involves severe brain pathology (Kirby et al., 1999), as in the case of Leigh syndrome (Rahman et al., 1996). Less profound, yet unexplained complex I defects appear to contribute to idiopathic Parkinson’s disease (PD) (Schapira et al., 1990; Keeney et al., 2006; Gatt et al., 2016; Flones et al., 2018) and likely other neurodegenerative disorders (Swerdlow, 2020; Terada et al., 2021). Most notably, pharmacological inhibition of complex I by exogenously applied toxins can evoke a Parkinson-like syndrome in animals and man (Langston et al., 1983; Sherer et al., 2007; Langston, 2017; Zeng et al., 2018) that in some models authentically recapitulates idiopathic PD (Betarbet et al., 2000; Cannon et al., 2009). Genetic mouse models of PD have been far from attaining the same degree of authenticity (Dawson et al., 2010; Creed and Goldberg, 2018; Chia et al., 2020). Among the complex I inhibitor-based models of PD, the MPTP/MPP^+^ model is the oldest (Langston et al., 1983; Langston, 2017) and arguably still the most widely employed model (Moosmann and Behl, 2002; Gibrat et al., 2009). Its dopaminergic cytotoxicity is essentially attributable to oxidative stress caused by complex I inhibition (Johannessen et al., 1986; Ramsay et al., 1987; Sayre et al., 2008; Hajieva et al., 2009).

Apparently unrelated to these lines of research, various epigenetic changes have been described to occur in PD. Specifically, decreased levels of global DNA cytosine methylation have been observed in post mortem brains from patients with PD and the related entity, dementia with Lewy bodies (Desplats et al., 2011). The effect was confirmed in CpG islands of regulatory regions of several disease-relevant genes, including the promotor and the first intron of α-synuclein (Jowaed et al., 2010; Matsumoto et al., 2010; Desplats et al., 2011) and in a number of other, potentially disease-relevant genes such as CYP2E1 (Kaut et al., 2022; Schaffner and Kobor, 2022). Characteristically altered, mostly decreased DNA methylation may also occur in blood cells from PD patients, suggesting a systemic phenomenon (Masliah et al., 2013).

Increased histone lysine acetylation is another notable epigenetic alteration in idiopathic PD. For instance, significant increases in H3K14 and H3K18 acetylation have been observed for the motor cortex of PD patients (Gebremedhin and Rademacher, 2016). These increases were yet contrasted by a decrease in H3K9 acetylation (Gebremedhin and Rademacher, 2016), which has been confirmed for the substantia nigra of unrelated PD cases (Harrison et al., 2018). Several other lysines have been found to be hyperacetylated in midbrain tissue (Park et al., 2016) and in the cerebral cortex (Toker et al., 2021) of idiopathic PD patients, the most prominent of which were H3K27 and, again, H3K14 (Toker et al., 2021). In summary, various histone lysines seem to be prone to hyperacetylation in PD. However, there is no consensus whether increased histone acetylation and the thereby induced transcriptional facilitation is adverse-pathologic (Park et al., 2016; Toker et al., 2021) or rather protective-adaptive (Kidd and Schneider, 2010) in the disease.

A connection between the two aforementioned signature elements of PD, complex I inhibition and epigenetic (dys)regulation, has been suggested by a small number of pivotal studies that have evidenced altered DNA methylation and histone acetylation in the wake of pharmacological complex I inhibition (Feng et al., 2015; Park et al., 2016; Yang et al., 2017). However, the purposefulness and origin of the potentially adverse (Matsumoto et al., 2010; Baeken et al., 2020; Toker et al., 2021) epigenetic transcriptional facilitation in PD has remained elusive. Hence, we have analyzed in more detail the transcriptomic events in human neuronal dopaminergic LUHMES cells after complex I inhibition. We find that mitochondria challenged in this way emanate a redox signal that is responsible for the selective transcriptional upregulation of mitochondrially imported gene products, particularly respiratory chain complex (RCC) subunits. We further characterize the mechanism of adaptive upregulation as partially epigenetic and related to redox-reversible DNMT3B and SIRT inhibition.

## Results

### Complex I inhibition causes widespread induction of nuclear-encoded RCC subunits in the absence of ATP depletion

The compound MPP^+^ (1-methyl-4-phenylpyridinium) is a frequently used reference tool for eliciting experimental complex I deficiency (Beal., 2001; Richardson et al., 2005; Langston, 2017). To ensure selectivity and avoid toxicity of this drug, differentiated human LUHMES cells were treated with 10 µM MPP^+^ for 48 h, following closely related protocols (Krug et al., 2014; Smirnova et al., 2016). No overt signs of toxicity were induced by this treatment regimen as reported (Krug et al., 2014; Baeken et al., 2020; Baeken et al., 2021), however, a moderate degree of microtubular reorganization was visible using immunocytochemistry (Suppl. Fig. 1). Transcriptomic analysis of the MPP^+^ treated cultures indicated that the majority of RCC subunits were transcriptionally induced by complex I inhibition, consistent with a functional, compensatory response (Fig. 1A; Suppl. Tab. 1). Specifically, 28 out of 37 complex I genes were significantly upregulated (Fig. 1C), as were 9 out of 10 complex III genes (Fig. 1E), 11 out of 11 complex IV genes (Fig. 1F), and 14 out of 16 complex V genes (Fig. 1G). On average, global transcription of complex I genes was induced by 61%, complex III genes by 106%, complex IV genes by 123%, and complex V genes by 76%.

**Figure 1.**
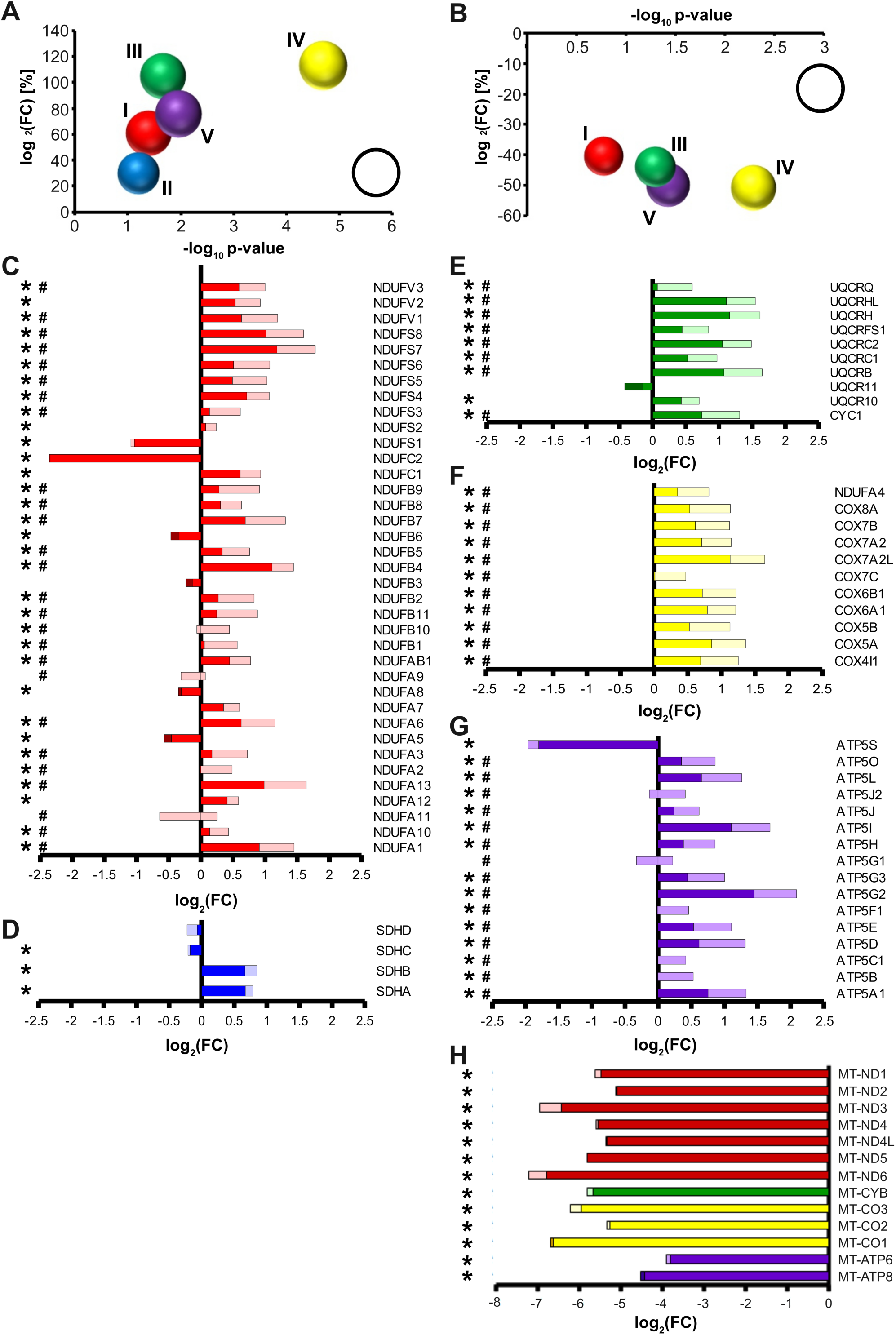
Redox-dependent transcriptional induction of nuclear-encoded RCC subunits following complex I inhibition. Transcriptional changes were measured by RNA sequencing of differentiated LUHMES cells treated with 10 µM MPP^+^ (a complex I inhibitor) and 20 nM PHT (a mitochondrial antioxidant) for 48h. Bubble diagram summarizing the regulation of RCC subunits after MPP^+^ treatment. Each bubble represents all nuclear subunits of one RCC (I - IV). The bubble position on the Y-axis indicates the log_2_ mean of the transcriptional fold changes (FC), the bubble position on the X-axis indicates the −log_10_ mean of the associated significance p values (n = 3, one-way ANOVA). The bubble size indicates the fraction of significantly regulated transcripts related to all transcripts (white bubble = 100%). **A.** Corresponding bubble diagram summarizing the effect of PHT on MPP^+^-treated cells. **B.** Individual regulation of nuclear-encoded complex I subunits. Regulation after MPP^+^ is indicated by total bar length; regulation after PHT/MPP^+^ is indicated by color coding: lighter color denotes that the MPP^+^ effect was reduced by PHT, darker color denotes that the MPP^+^ effect was increased by PHT. Symbols indicate: *p-value ≤ 0.05 for MPP^+^ vs. control, ^#^p-value ≤ 0.05 for PHT/MPP^+^ vs. MPP^+^ by one-way ANOVA (n = 3) in all panels of this figure. **C.** Individual regulation of complex II subunits. **D.** Individual regulation of nuclear-encoded complex III subunits. **E.** Individual regulation of nuclear-encoded complex IV subunits. **F.** Individual regulation of nuclear-encoded complex V subunits. **G.** Regulation of mitochondrially encoded RCC subunits.

In inhibiting complex I electron flow from the aqueous NADH oxidation site to the ubiquinone binding site, MPP^+^ causes two major biochemical effects: the loss of complex I as a proton pump contributing to ATP generation, and the production of superoxide radicals and other reactive oxygen species (ROS) by complex I (Fallon et al., 1997; Beal., 2001; Richardson et al., 2005). To distinguish which of these effects caused the transcriptional changes, we applied the mitochondrial antioxidant phenothiazine (PHT) to MPP^+^ treated cells. PHT is a nanomolar-activity antioxidant compound that permeates mitochondria and has shown high efficacy against mitochondrial ROS even in models where classic phenolic antioxidants generally fail (Hajieva et al., 2009; Mocko et al., 2010; Tapias et al., 2019). However, PHT is not a two-electron reductant, meaning that it does not shuttle electrons from inhibited complex I to complex IV such as methylene blue (Atamna et al., 2008; Ohlow and Moosmann, 2011). Thus, it cannot ameliorate bioenergetic deficits, but merely acts as a selective ROS scavenger.

PHT treatment caused a significant, apparently uniform blunting of the transcriptional effects of MPP^+^ (Fig. 1B; Suppl. Tab. 1). More specifically, 26 out of 37 complex I genes were significantly downregulated compared to MPP^+^ only-treated cells (Fig. 1C), as were 8 out of 10 complex III genes (Fig. 1E), 11 out of 11 complex IV genes (Fig. 1F), and 15 out of 16 complex V genes (Fig. 1G). Across all subunits, transcription of complex I genes in these cells was reduced by 40%, complex III genes by 44%, complex IV genes by 51%, and complex V genes by 50%. Hence, approximately half of the regulatory effect of complex I inhibition was prevented by a low nanomolar dose of PHT despite the inherent kinetic imperfection of any radical scavenging system.

Notably, all mitochondrially encoded transcripts were severely reduced following MPP^+^ treatment (Fig. 1H), consistent with earlier reports (Krug et al., 2014). Most likely, the engagement of the cells in preparatory mitochondrial DNA replication, which is known to be incompatible with mitochondrial transcription (Agaronyan et al., 2015), accounts for this effect. Mitochondrial transcripts were also not responsive to PHT treatment like those nuclear encoded RCC subunits that had been suppressed by MPP^+^ treatment, indicating their disparate, redox-independent regulation. Interestingly, the three most prominently suppressed nuclear encoded RCC-related genes were either supernumerary, regulatory subunits involved in RCC assembly, like ATP5S (Belogrudov, 2009) and NDUFC2 (Alahmad et al., 2020), or regulate supercomplex formation and have additional functions in the cytosol, like NDUFS1 (Elkholi et al., 2019). Plausibly, assembly factors are less required as long as mitochondrial transcription and translation do not proceed.

To experimentally ascertain the supposed non-interference of PHT with cellular bioenergetics under the employed conditions, LUHMES cells treated identically as before were surveyed for a series of functional metabolic readouts. As shown in Fig. 2, MPP^+^ treatment had no significant effect on cellular ATP levels, without or with PHT co-administration (Fig. 2A). NAD^+^ levels were also unchanged (Fig. 2B), while NADH levels were increased by complex I inhibition as expected (Fig. 2C), resulting in a significant drop in the NAD^+^/NADH ratio relevant for metabolic flux (Fig. 2D). PHT had no modulatory effect on any of these parameters. Moreover, PHT did not alter the expectable effects of MPP^+^ on lactate (Fig. 2E) and glucose (Fig. 2F), respectively. In contrast, analysis of cellular ROS levels by means of an indicator dye, CellROX, clearly demonstrated the prooxidant effect of MPP^+^ and the antioxidant effect of PHT (Fig. 2G and 2H), as widely reported (Beal., 2001; Hajieva et al., 2009; Mocko et al., 2010; Tapias et al., 2019). Finally, assessment of cell culture pH indicated a moderate acidification in the wake of MPP^+^ treatment as expected (Fig. 2I), accompanied by carboanhydrase induction (Fig. 2J). PHT, in turn, did not significantly modulate the pH change and had an only minor effect on 2 out of 5 of the detected carboanhydrases. In summary, PHT was incapable of modulating the bioenergetic effects of complex I inhibition, but selectively blunted the ROS effect. Besides, it had a small effect on the NAD^+^/NADH ratio at baseline (Fig. 2D).

**Figure 2.**
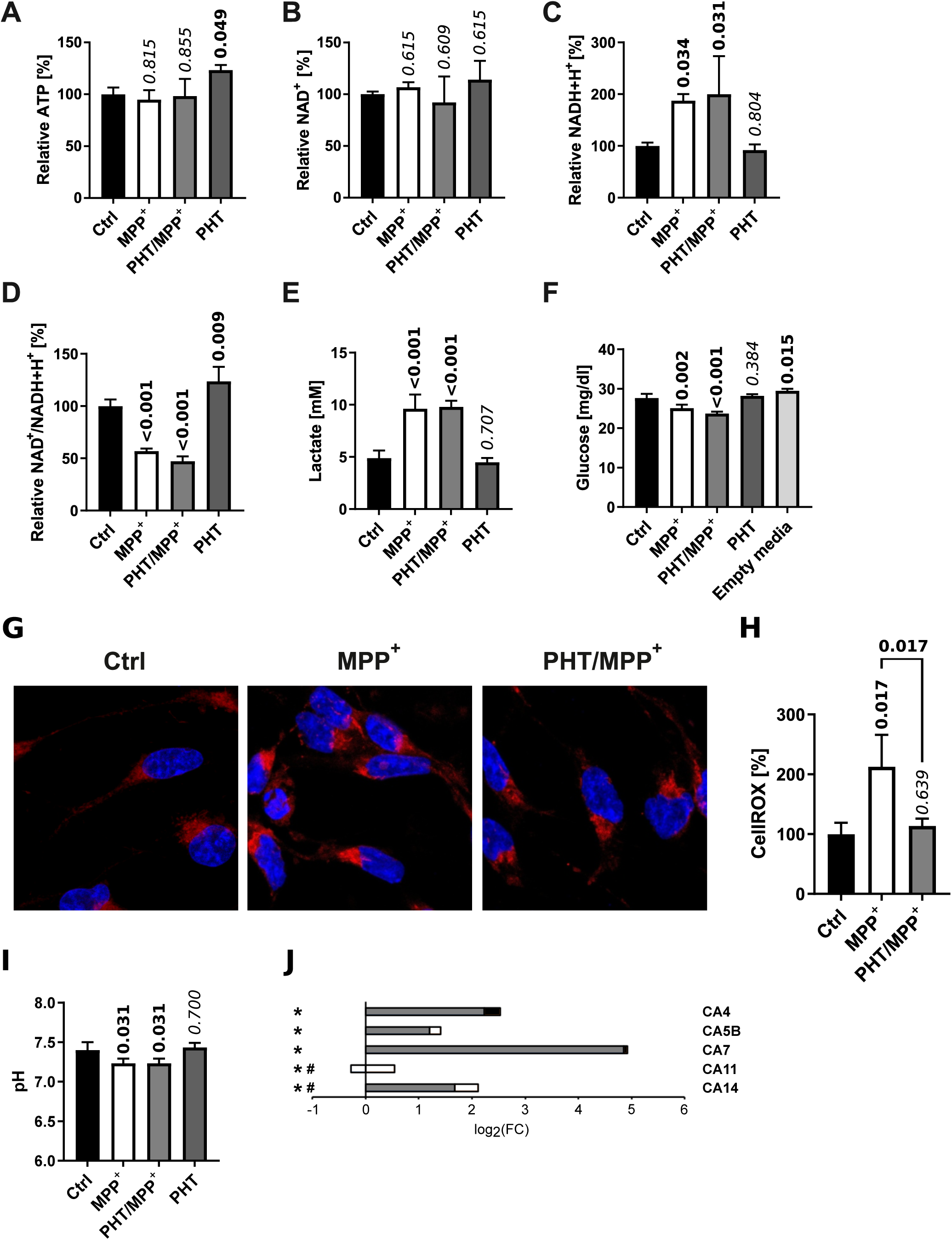
Bioenergetic and redox changes evoked by complex I inhibition. **A.** Relative ATP levels in cells treated with 10 µM MPP^+^ and 20 nM PHT for 48 h identically as in the experiment in Fig. 1. Numbers atop of the bars indicate the level of significance versus the control as determined by two-way ANOVA and Benjamini-Hochberg post-hoc test. Bold numericals highlight p ≤ 0.05. All standard deviations in this figure result from n = 3 experiments. **B.** Relative levels of NAD^+^ in cells treated as before. **C.** Relative levels of NADH+H^+^ in the same cells. **D.** NAD^+^/NADH+H^+^ ratio in the same cells. **E.** Lactate concentrations in 48 h culture media from the same cells. **E.** Glucose concentrations in 48 h culture media from the same cells. Fresh media were assayed for control purposes. **G.** Fluorescence images of LUHMES cells administered with the ROS indicator compound CellROX (red) and the chromatin stain Hoechst 33258 (blue). **H.** Densitometric quantification of CellROX fluorescence intensity normalized to the number of cells assayed. **I.** Level of acidification in 48 h culture media from cells treated as before. **J.** Transcriptional regulation of all five carboanhydrase enzymes detected in LUHMES cells by RNA sequencing. Symbols indicate: *p-value ≤ 0.05 for MPP^+^ vs. control, ^#^p-value ≤ 0.05 for PHT/MPP^+^ vs MPP^+^. Shading of the fill color is used to denote PHT effects as in Fig. 1.

### MPP^+^ causes global DNA hypomethylation through DNMT3B insufficiency that is responsive to the antioxidant PHT in vitro

Considering the rather uniform up- and down-regulation of many different genes at disparate loci (Suppl. Fig. 2) in the nucleus, we hypothesized that epigenetic mechanisms might be critically involved. Indeed, decreased levels of DNA methylation have been reported in different models of PD and in patient-derived tissue (Desplats et al., 2011; Yang et al., 2017). Consistently, MPP^+^ caused a substantial reduction of global 5-methylcytosine levels in the nucleus of differentiated LUHMES cells within 48 h (Fig. 3A,B). This effect was abrogated by PHT cotreatment. The control drug 6-thioguanine, an established DNA methylation suppressor (Agrawal et al., 2018), elicited comparable effects at the employed standard concentration of 1 µM (Mender et al., 2015).

**Figure 3.**
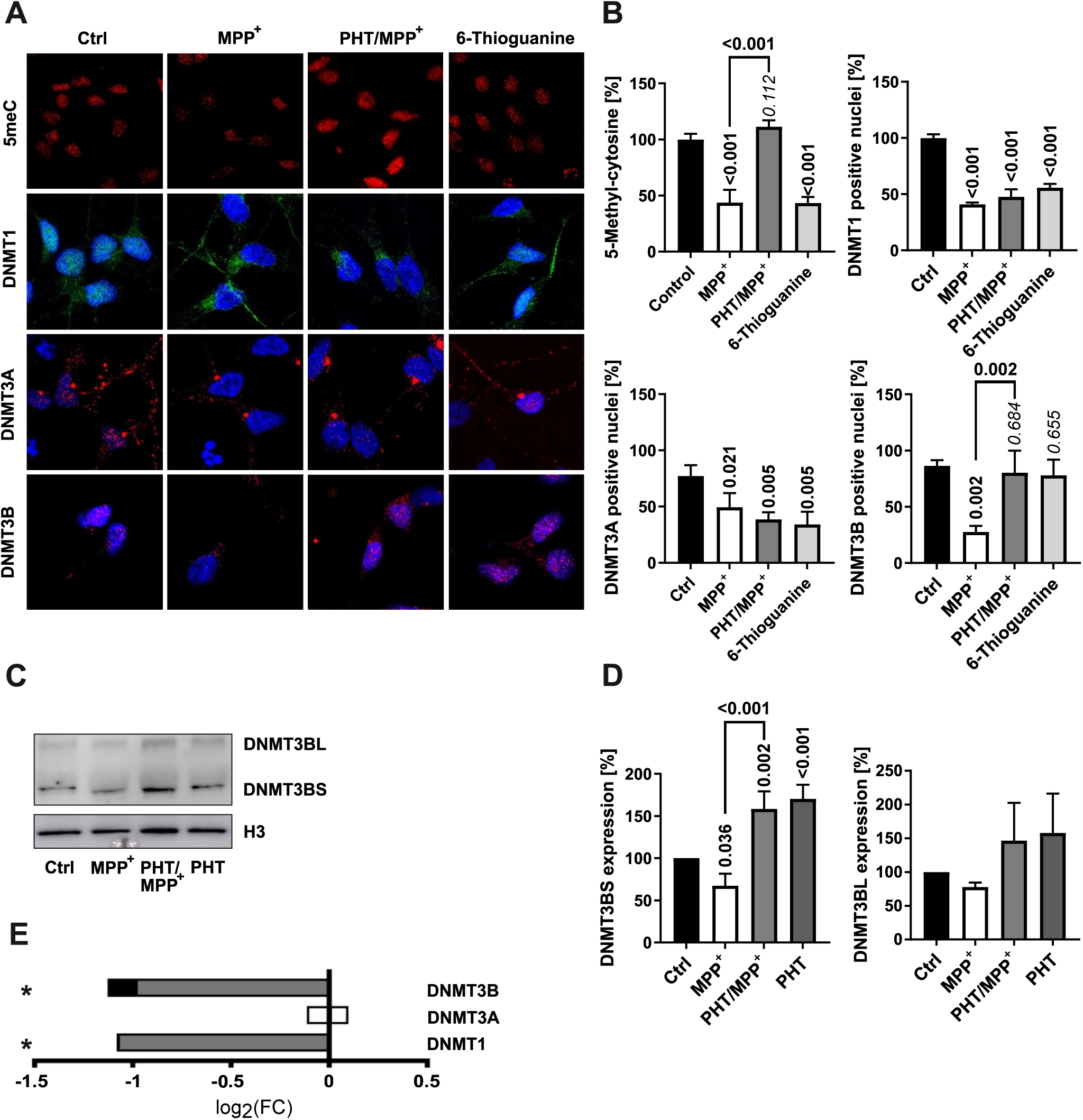
Changes in DNA methylation and DNMT localization. **A.** Fluorescence microscopic images of immunostained LUHMES cells treated with 10 µM MPP^+^, 20 nM PHT, or 1 µM 6-thioguanine over 48 h (63x magnification). In the first row, 5-methylcytosine (5meC) is visualized in red. The other rows depict DNMT1 (green), DNMT3A (red), and DNMT3B (red) as indicated. The blue staining represents the chromatin dye Hoechst 33258. **B.** Bar graph diagrams of densitometric quantifications of images (n = 3, number of cells per n: 50-100) as shown in A. Numbers atop of the bars and bold numericals (p ≤ 0.05) are used as in Fig. 2.. **C.** Western blotting of DNMT3B and H3 in cells treated with 10 µM MPP^+^ and 20 nM PHT as before. DNMT3B features two bands at ∼100 kDa (DNMT3BL) and ∼70 kDa (DNMT3BS). **D.** Densitometric quantification of n = 3 DNMT3B blots normalized on H3. **E.** Transcriptional regulation of the DNMTs. Symbols indicate: *p-value ≤ 0.05 for MPP^+^ vs. control. Shading of the fill color is used to denote PHT effects as in Fig. 1.

DNA methylation in humans is established by three DNA methyltransferases. DNMT1 is considered a maintenance protein requiring a hemi-methylated template, whereas DNMT3A and DNMT3B are de novo methyltransfereases (Hermann et al., 2004). Analysis of DNMT1 and DNMT3B by immunocytochemistry indicated that these proteins were mainly localized to the nucleus whereas DNMT3A localized to the nucleus and the cytosol, but experienced perinuclear condensation under MPP^+^ administration (Fig. 3A,B). All three proteins exhibited a significant shift towards the cytosol upon MPP^+^ exposure (Fig. 3A,B), as also reported for DNMT1 in PD (Desplats et al., 2011). Notably, these effects were not rescued by PHT cotreatment except in the case of DNMT3B, which was therefore putatively assigned responsible for the observed changes in DNA methylation. The compound 6-thioguanine had a similar effect on the localization of DNMT1 and DNMT3A as MPP^+^, but did not influence DNMT3B. The former observation is consistent with the reported induction of proteasomal degradation of DNMT1 by 6-thioguanine (Yuan et al., 2011), which may also affect DNMT3A (Huang et al., 2022), but potentially spares DNMT3B.

DNMT3B expression in the Western blot showed two bands at ∼95 kDa (DNMT3BL) and ∼72kDa (DNMT3BS) (Fig. 3C,D). MPP^+^ caused a modest decline of both isoforms by approximately 30%, which was prevented by PHT treatment, but significantly only for DNMT3BS, the major form of DNMT3B in these cells. Exclusive PHT treatment also showed significantly increased protein levels of DNMT3BS compared to the control group (Fig. 3C,D) without rescuing DNMT3B at the mRNA level (Fig. 3E). These data suggest that DNMT3B is inducibly degraded in the cytosol in response to a redox signal that can be suppressed by PHT, which leads to lowered DNMT3B activity in the nucleus upon early mitochondrial distress. The baseline suppression of DNMT1 and DNMT3B transcription by MPP^+^, which was PHT-indifferent (Fig. 3E), arguably relates to another biochemical mechanism, but may indeed amplify the efficacy of the redox signal. Like other epigenetic effector proteins (Park et al., 2016; Baeken et al., 2021; Baeken, 2023), DNMTs are now well established to be functionally regulated by their controlled proteolytic degradation (Du et al., 2010; Yuan et al., 2011; Huang et al., 2022). DNMT3A/3B have also been shown to be affected by the thiol redox state of the cell (Chen et al., 2012).

### MPP^+^ causes global lysine hyperacetylation through SIRT1 suppression that is responsive to the antioxidant PHT in vitro

Histone lysine acetylation appears to be widely induced in idiopathic PD (Gebremedhin and Rademacher, 2016; Park et al., 2016; Toker et al., 2021) and in models of PD based on complex I inhibition (Park et al., 2016). Of the many sites that appear to be hyperacetylated in PD, including H2K15, H3K14, H3K18, H3K27 and H4K5, the site H3K14 may represent one of the most reproducible disease markers. No conflicting data as for H3K9 have been reported for H3K14 (Gebremedhin and Rademacher, 2016; Park et al., 2016), and it may also be more stringently induced than the otherwise related, more widely explored H3K9 site (Karmodiya et al., 2012). H3K14 has not been investigated after complex I inhibition.

Treatment of LUHMES with MPP^+^ as before evoked a significant increase in total lysine acetylation as well as H3K14 acetylation, which were entirely prevented by PHT treatment and thus redox-related (Fig. 4A-D). The increase in H3K14 acetylation as per Western blot (∼600%) (Fig. 4C) vastly exceeded the increase in total lysine acetylation (∼30%) (Fig. 4D). Two minimally modified PHT derivatives that essentially lack antioxidant activity (Hajieva et al., 2009), namely N-methylphenothiazine (MPHT) and N-acetylphenothiazine (APHT), were also tested in this assay because putatively involved histone deacetylases of the sirtuin family are highly sensitive to the levels of the somewhat related heteroaromatic molecule NAD^+^, containing nicotinamide (Anderson et al., 2017). MPHT and APHT were clearly less potent than PHT in their prevention of lysine acetylation (Fig. 4C,D), corroborating that the PHT effect was caused by antioxidation.

**Figure 4.**
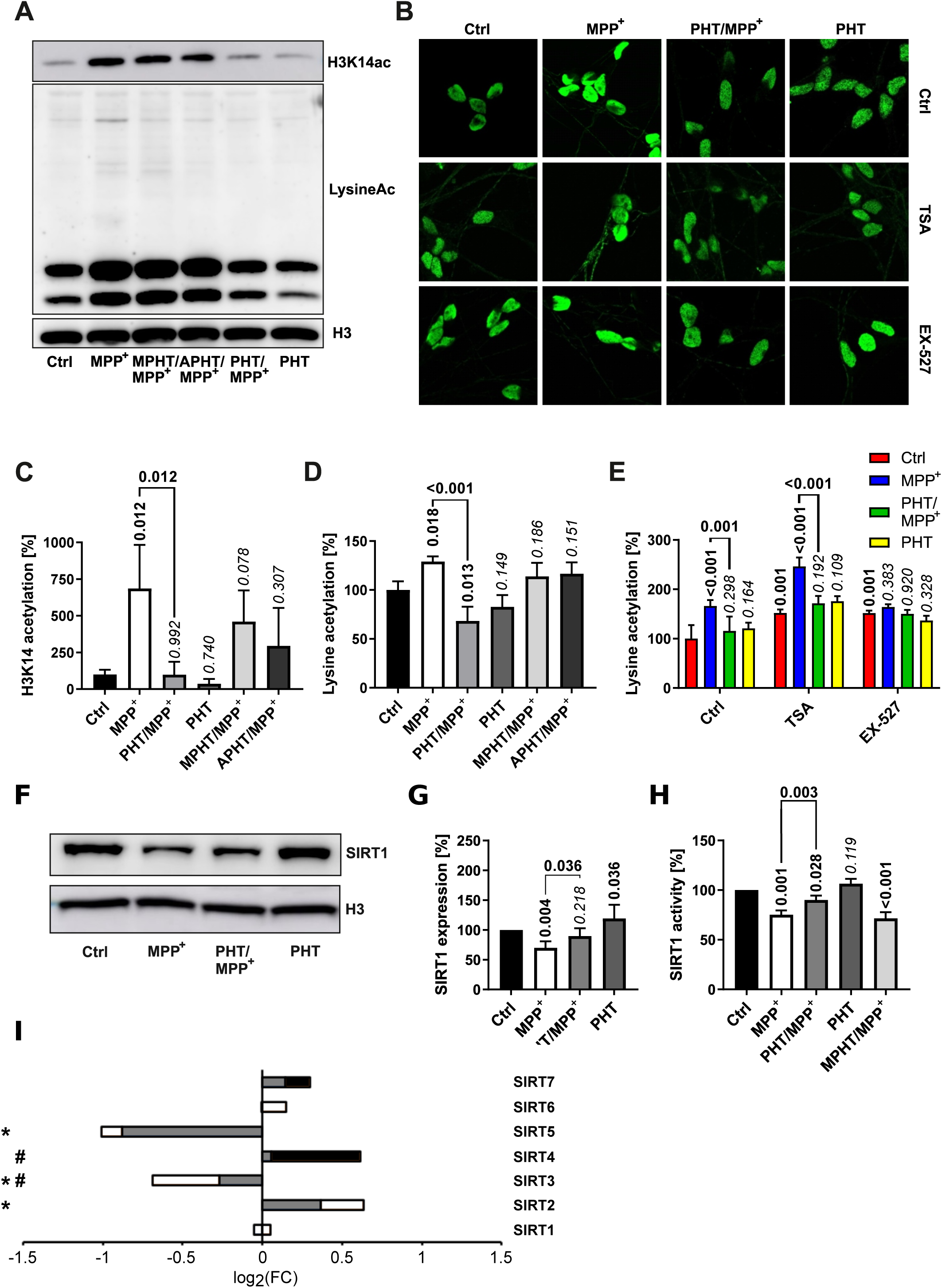
Lysine acetylation and deacetylase expression. Lysine acetylation and the expression of selected deacetylases was investigated in LUHMES cells treated with 10 µM MPP^+^ and 20 nM PHT for 48 h as before. MPHT and APHT are two inactive PHT congeners used at 20 nM concentration as PHT. **A.** Western blotting against H3K14ac, total acetylated lysine and H3. **B.** Microscopic images (63x magnification) of cells immunostained for total lysine acetylation. TSA (50 nM) and Ex-527 (100 nM) are (class-)specific deacetylase inhibitors. **C.** Densitometric quantification of Western blots (n = 3) against H3K14ac as shown in A, normalized on H3. **D.** The same quantification (n = 3) done for total lysine acetylation. **E.** Quantification of lysine acetylation by image analysis of immunostained cells as shown in B (n = 3, number of cells per n: 50-100). **F.** Western blotting of the NAD-dependent deacetylase SIRT1 and H3. **G.** Densitometric quantification of n = 3 SIRT1 blots normalized on H3. **H.** SIRT1 deacetylase activity determined in lysates of differentiated LUHMES cells (n = 3). **I.** Transcriptional regulation of all sirtuins detected in LUHMES cells by RNA sequencing. Symbols indicate: *p-value ≤ 0.05 for MPP^+^ vs. control, #p-value ≤ 0.05 for PHT/MPP^+^ vs MPP^+^. Shading of the fill color is used to denote PHT effects as in Fig. 1.

To further define the cause of MPP^+^-induced hyperacetylation, we employed TSA, an inhibitor of Zn^2+^-dependent HDACs, and EX-527, a SIRT1 inhibitor with approximately 200-fold selectivity against SIRT2 and SIRT3 (Yoshida et al., 1999; Napper et al., 2005). Analysis of global acetylation by immunocytochemistry (Fig. 4B) and normalized image quantification (Fig. 4E) indicated that both agents caused an increase in acetylation by approximately 50% and, thus, resembled MPP^+^ quantitatively. However, these increases were resistant to antioxidant PHT treatment as expected. Additive treatment with the inhibitors plus MPP^+^ gave a further (and PHT-reversible) increase only with TSA, but not with EX-527 (Fig. 4E). Thus, the effect of MPP^+^ was concluded to be mediated by EX-527-inhibited deacetylases, but not by TSA-inhibited deacetylases.

Investigation of SIRT1 by Western blot (Fig. 4F,G) indicated that this protein was significantly reduced upon MPP^+^-treatment in a PHT-reversible fashion; an according result was obtained in a direct, fluorescent SIRT1 activity assay with the cell lysates (Fig. 4H). Here, the enzyme activity loss in the lysate was partially prevented by PHT, but unaltered by MPHT. In view of the unchanged transcription of SIRT1 in these cells (Fig. 4I), these results demonstrate a loss of the protein SIRT1 and its enzyme activity due to a redox signal induced by mitochondrial complex I inhibition. The loss of this protein is likely attributable to the redox-dependent induction of autophagic degradation of most sirtuins by MPP^+^ as recently reported (Baeken et al., 2021). Because SIRT1 enzyme activity is also known to be negatively affected by direct cysteine oxidation (Shao et al., 2014; Kalous et al., 2020), it is possible that both mechanisms operate in parallel.

### MPTP causes ROS-dependent epigenetic changes in vivo

Redox biological experiments in cell culture involve the general danger of returning exaggerated effects due to an unphysiologically oxidative environment (Kunath et al., 2020). Hence, we have tested the validity of some of the described molecular events in a mouse model of PD based on the same initiating event, namely complex I inhibition by MPP^+^. Therefore, the pro-toxin MPTP (Beal., 2001; Langston, 2017) was administered intraperitoneally to 10 week-old male C57Bl/6J mice as sketched in the scheme in Fig. 5. PHT was administered orally, including a roll-in period (for details, compare the Materials and Methods). Doses were chosen as to obtain an intermediate degree of toxicity only in the particularly vulnerable region substantia nigra (SN), as widespread cell death would arguably give rise to strong secondary effects.

**Figure 5.**
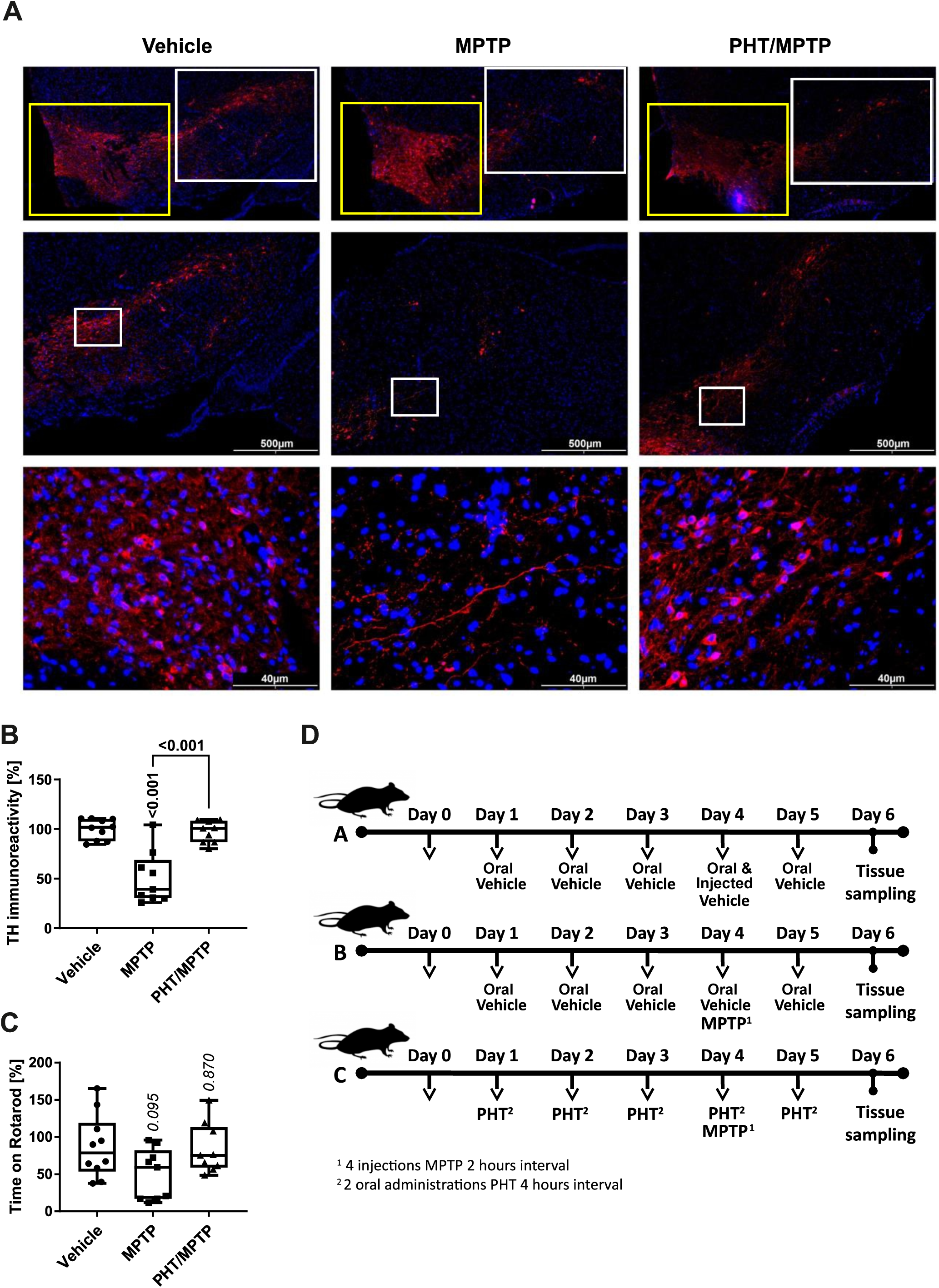
General characterization of the MPP^+^/PHT system in vivo. Male, wild-type, 10-week old C57Bl/6J mice were treated with the metabolic MPP^+^-precursor MPTP and analyzed behaviorally and biochemically. **A.** Fluorescence immunohistochemistry of midbrain slices stained for the dopaminergic marker tyrosine hydroxylase (TH, red), counterstained with Hoechst 33258 (blue). The upper row shows an overview of the rostral midbrain depicting the VTA (ventral tegmental area; yellow box) on the left, and the SN (substantia nigra; white box) on the right. The middle row displays 4x magnifications of the SN as approximated in the upper row. The lower row shows 20x magnifications of the central SN. **B.** Quantification of TH immunoreactivity in sections from all treatment groups (n = 9-10 animals, each dot representing one animal). **C.** Rotarod performance of the different animals (n = 9-10 animals). Rotarod is a widely employed motor diagnostic test in the study of parkinsonism. **D.** Schematic overview of the different treatment groups. Group A received two types of vehicles (oral vehicle and injected vehicle), group B received oral vehicle and MPTP, group C received PHT and MPTP (including a PHT roll-in period) as displayed. Note that PHT was administered orally (DMSO:corn oil 1:50), whereas MPTP was injected intraperitoneally (0.9% saline).

To verify the efficacy and selectivity of the employed MPTP dose and to probe any potential prevention by PHT, we analyzed the expression of tyrosine hydroxylase (TH), a commonly adopted marker of dopaminergic cell viability in vivo (Beal., 2001; Langston, 2017). Hence, midbrain slices were immunostained with antibodies against TH and counterstained with the chromatin dye Hoechst 33258 (Fig. 5A). The staining revealed a visible drop in the number of TH-positive cells in the SN, the primary region affected in PD (Fig. 5A). The adjacent, much larger ventral tegmental area (VTA) were spared from toxicity as expected (Beal, 2001). Quantification by image analysis involving cell-cell border demarcation and counting of TH-positive cells yielded approximately 50% loss of SN neurons, with PHT treatment affording almost complete protection (Fig. 5B) as described earlier in related models (Mocko et al., 2010; Tapias et al., 2019). Motor performance experiments (“Rotarod”) done with the animals before sacrifice suggested a variable decline of capabilities from MPTP treatment including rescue by PHT, but statistical significance was not reached (Fig. 5C).

Analysis of H3K14 acetylation in the SN demonstrated that MPTP treatment triggered this modification in vivo in a PHT-reversible fashion, with an increase of about 50% as per immunocytochemistry (Fig. 6A,C) and about 150% as per Western blot (Fig. 6B,D). Global lysine acetylation as per Western blot was raised by about 50% (Fig. 6E). These results recapitulated the human cell culture outcome (Fig. 4), but to an attenuated degree. However, SIRT1 expression in the mouse in vivo was not decreased, but rather increased (Fig. 6F). Since PHT did not revert this effect, a redox-unrelated mechanism may have dominated here. Two additional SIRTs known to be rapidly degraded in vitro after MPP^+^ treatment were probed for control purposes, and indeed, SIRT3 and SIRT4 were suppressed by about 30% and 20%, respectively, and PHT-reversibly in vivo (Fig. 6G,H). The origin of the differential behavior of SIRT1 is unclear at present. DNMT3B expression, in turn, was decreased in vivo by about 30% and rescued by PHT administration, which recapitulated the in vitro situation. In summary, these findings confirm the operability of a redox signal targeting epigenetic regulator proteins after complex I inhibition in vivo.

**Figure 6.**
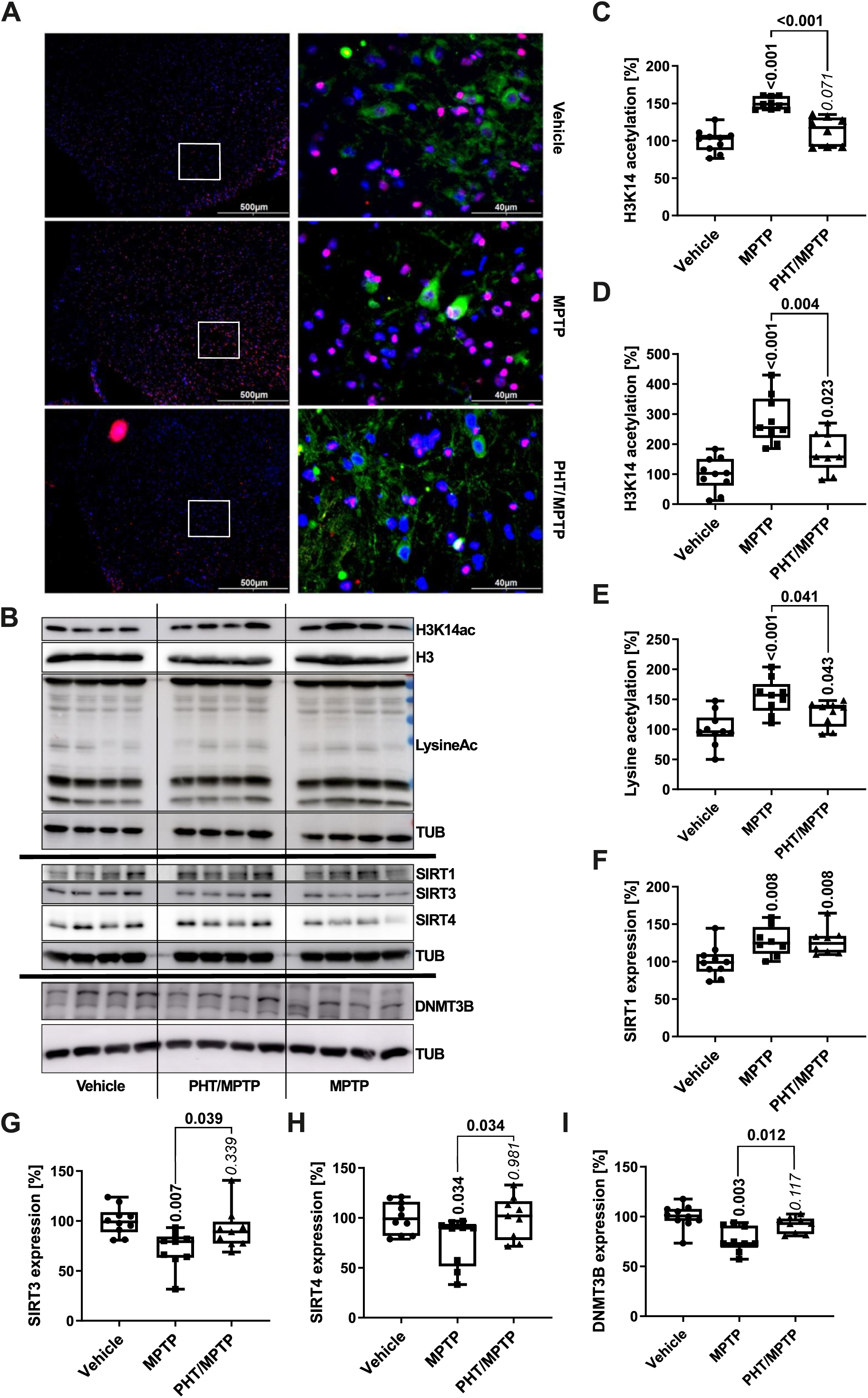
Modulation of epigenetic markers by complex I inhibition in vivo. **A.** Immunomicrographs of mouse midbrain SN slices stained for H3K14ac (red) and the dopamine transporter DAT (green), counterstained with Hoechst 33258 (blue). Left column, 4x magnification, right column, 20x magnification. **B.** Representative Western blots of midbrain lysates (4 mice per group) for H3K14ac, H3, total lysine acetylation, α-tubulin (TUB), SIRT1, SIRT3, SIRT4 and DNMT3B. **C.** Image analytical quantification of the H3K14ac signal from immunohistochemistry as in A, normalized to total nuclei (n = 3 slices from 9-10 animals, number of cells per n: approximately 1000). **D.** Densitometric quantification of the H3K14ac signal from Western blotting as in B, normalized to H3 expression (n = 9-10 animals). **E.** The same analysis as in D for total lysine acetylation, normalized to TUB expression (n = 9-10). **F.-I.** Analogous Western blot quantifications of SIRT1, SIRT3, SIRT4 and DNMT3B, all normalized to TUB (n = 9-10).

### The transcriptomic effects of the MPP^+^-induced redox signal are specific for mitochondrial targets and purposefully related genes

From the results presented so far, the hypothesis was derived that the redox-dependent effect of complex I inhibition towards the epigenetic switches SIRT and DNMT3B may be part of a targeted regulatory cycle to specifically maintain mitochondrial metabolic functioning. This hypothesis was tested by revisiting the transcriptomic changes that accompanied the induction of most RCC subunits already described in Figure 1. Out of 15168 genes recovered in the RNA sequencing experiment, of which 3639 were upregulated, and 3531 were downregulated. A total of 826 mitochondrially imported genes (“mito-genes”) and 78 RCC complex genes were selected for statistical analysis; a p < 0.001 significance level was adopted to account for the increased number of comparisons.

Mito-genes were found to be significantly more highly induced than average genes. Of these mito-genes, RCC complex genes were significantly more often induced than the other genes (Fig. 7A). Hence, complex I inhibition caused a selective increase in the transcription of genes counteracting any potential malfunction of complex I and the respiratory chain. The selective targeting of mito-genes and RCC complex genes by MPP^+^ was also selectively affected by PHT treatment: co-application of PHT lowered mito-genes significantly more highly than average genes, and RCC complex genes significantly more often than other mito-genes (Fig. 7B). Notably, the effect of PHT was strong enough to entirely abrogate any group-specific targeting of complex I inhibition (Fig. 7C), suggesting that the statistical selectivity of the transcriptional response was largely mediated by redox signaling, and not by other mito-nuclear signaling mechanisms, several of which have been described (Quiros et al., 2016; Walker and Moraes, 2022).

**Figure 7.**
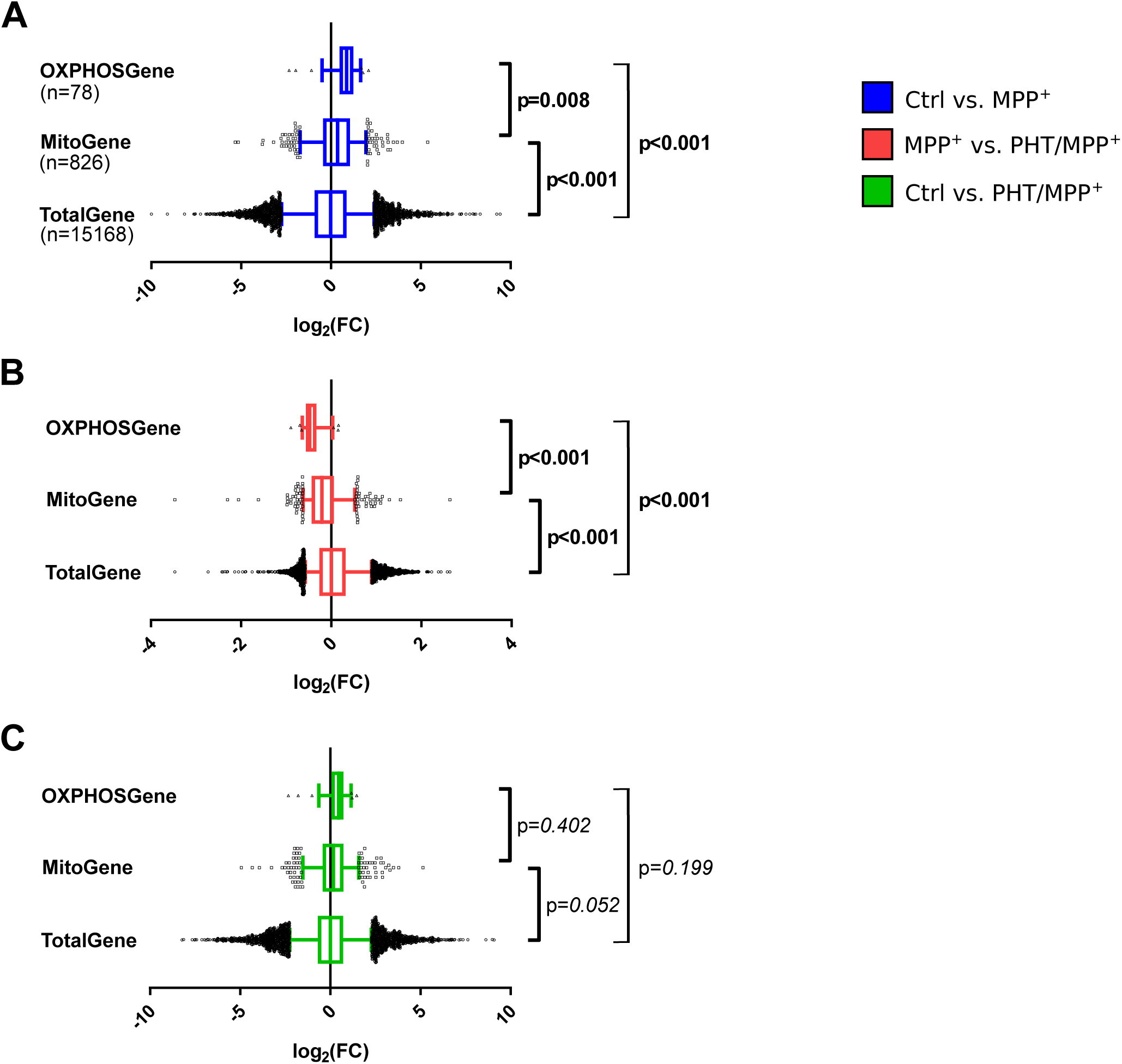
Selectivity of the transcriptomic changes induced by complex I inhibition for mitochondrial genes. Box plot analysis of gene expression patterns in differentiated LUHMES cells treated with 10 µM MPP^+^, 20 nM PHT, or both for 48 h as in Figure 1. A comprehensive selection of 78 RCC genes (OXPHOSGene) were compared with 826 mitochondrially imported genes (MitoGene) and the total, recovered transcriptome of 15168 genes (TotalGene). Statistical analysis was done by One-way ANOVA**A.** Comparison of MPP^+^-treated cells with control cells. **B.** Comparison of MPP^+^-treated cells with PHT- and MPP^+^-treated cells. **C.** Comparison of control cells with PHT- and MPP^+^-treated cells.

These results were compared with an arbitrary selection of other metabolic gene clusters including, for example, 93 autophagy-related genes and 28 nucleotide catabolism-related genes (Fig. 8). Of nine investigated clusters, two were significantly induced by MPP^+^ treatment, namely glycolysis and cholesterol biosynthesis (Fig. 8A). PHT treatment had a significant suppressive effect on both MPP^+^-induced clusters and, in addition, on the amino acid catabolism cluster (Fig. 8B). In the direct comparison of the control transcriptome with the MPP^+^/PHT transcriptome, none of the clusters attained significance. Closest to significance were, again, glycolysis and cholesterol biosynthesis as being modestly induced. These analyses demonstrate that redox signaling was responsible not only for the transcriptional targeting towards the mitochondrion, but also for the transcriptional targeting towards a functionally related metabolic pathway, glycolysis. The relationship between complex I inhibition and induced cholesterol biosynthesis is elusive, even if ample literature exists as regards cholesterol and PD (Garcia-Sanz et al., 2021).

**Figure 8.**
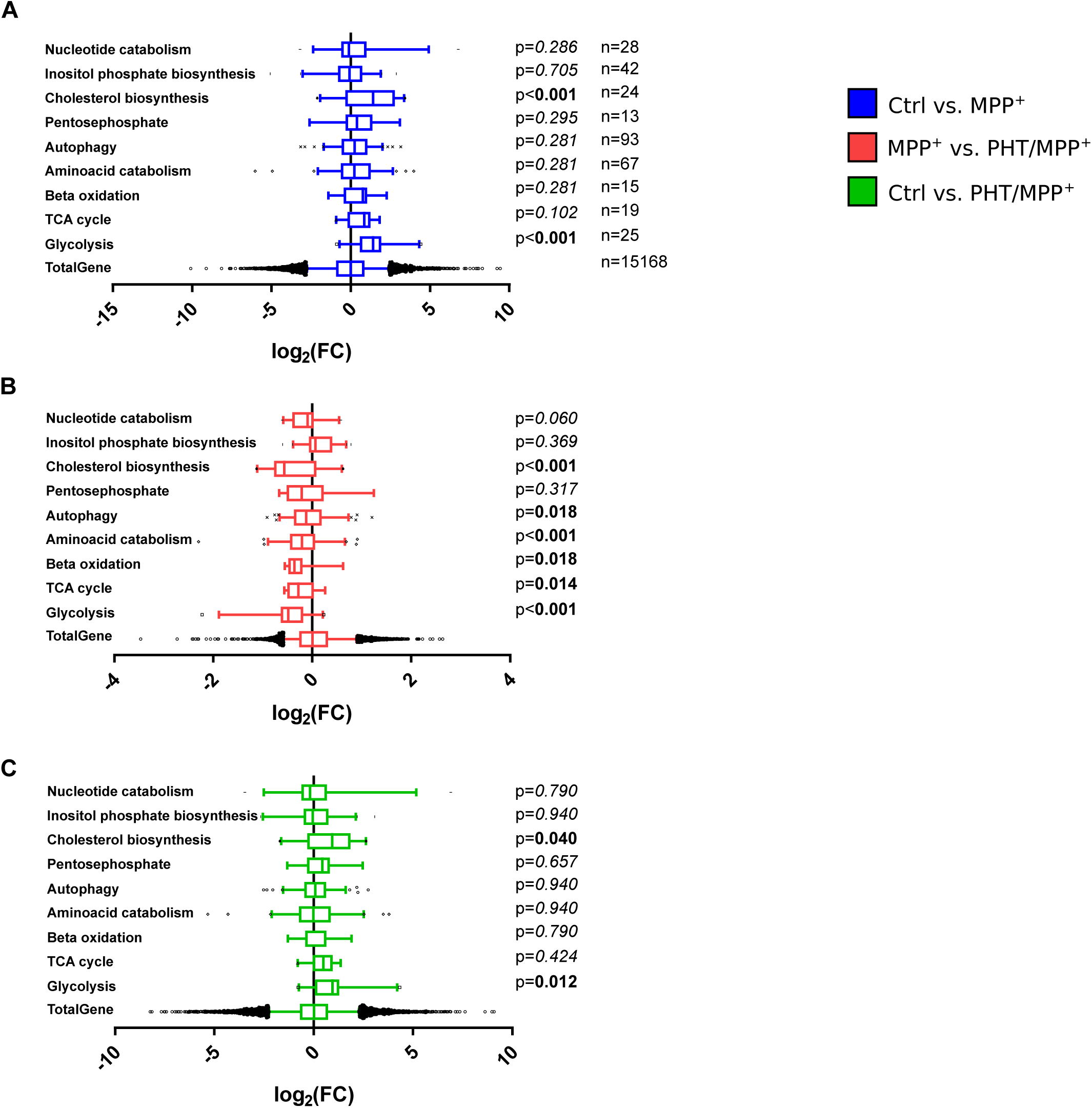
**Effects of complex I inhibition on general metabolic pathways.** Box plot analysis of gene expression patterns as in Figure 7. The indicated numbers of genes related to different metabolic pathways as per the Reactome database were statistically compared by One-way ANOVA. **A.** Comparison of MPP^+^-treated cells with control cells. **B.** Comparison of MPP^+^-treated cells with PHT- and MPP^+^-treated cells. **C.** Comparison of control cells with PHT- and MPP^+^-treated cells.

## Discussion

Complex I is the main point to entry of electrons into the respiratory chain (Lenaz and Genova, 2009). This also appears to apply to dopaminergic neuronal cells (Risiglione et al., 2020). Despite its overall high expression in the range of several million copies per cell (Wisniewski et al., 2014), complex I does not seem to be present in large excess, as can be judged from a series of observations. First, patients with severe mitochondrial disease due to complex I mutations still exhibit relatively high residual activities. Affected patients often present with 20-40% residual activity in muscle biopsies and 30-100% activity in fibroblasts (Rahman et al., 1996; Kirby et al., 1999; Loeffen et al., 2000; Bugiani et al., 2004), whereas 10-20% residual activity usually results in lethality (Kirby et al., 1999). Second, despite being the major ubiquinone reductase to fuel complex III and complex IV in a quasi-linear reaction, it is expressed at a much lower level than the other complexes; specifically, complexes I:III:IV are expressed at a ratio of approximately 1:3:6 in different tissues and species (Lenaz and Genova, 2009). Moreover, complex I downregulation has emerged as key adaptation of long-lived animals (Lambert et al., 2010; Munro et al., 2013; Mota-Martorell et al., 2020), suggesting that the exceptionally long-lived H. sapiens may also operate at the lowest possible complex I level to avoid reverse electron transport and, thereby, life-shortening ROS overproduction (Mookerjee et al., 2010; Mota-Martorell et al., 2020).

These observations indicate that any incidental losses of complex I activity need to be detected early by the cell, in order to induce compensatory measures, particularly a transcriptional induction of complex I and potentially other RCC subunits in the nucleus. Preferably, such losses should be detected before severe bioenergetic deficits such as ATP depletion accrue. In the present study, we show that this is indeed the case in human cells, and we demonstrate that the imminent danger signal is an oxidant that can be suppressed by low doses of a radical scavenger. Using an oxidant signal for this purpose appears to be particularly expedient because oxidant production by complex I and III sites is known to be induced rapidly and rather universally in response to diverse pharmacological agents that inhibit RCCs (Tahara et al., 2009; Brand, 2016; Wong et al., 2017). Intriguingly, complex I inhibitors with low toxicity but high anti-diabetic efficacy (i.e., metformin versus other bi-/diguanides) appear to be characterized by relatively modest suppression of primary catalysis combined with relatively high induction of superoxide production (Cameron et al., 2018). The latter would start metabolic transcriptional reprogramming according to the observations described herein.

The fact that multiple chemical inhibitors of RCCs cause superoxide production implies that multiple protein damaging events such as denaturation, cofactor loss or lipoxidation will likely evoke the same effect. Hence, impairment of any of the complexes I, III or IV by wear and tear will arguably entail increased superoxide production by either the damaged or an upstream complex. The induced superoxide response is therefore rather nonspecific because the precise site of damage is not conveyed to the nucleus. However, it is rapid and emanates from the mitochondrion in the wake of a still only functional and perhaps reversible, but not yet structural insufficiency. Both aspects are advantageous because firstly, there will still be enough ATP to afford protein biosynthesis and thus repair, and secondly, because extensive protein damage triggering an unfolded protein response (Zhao et al., 2002; Quiros et al., 2016) is not required to release the repair signal. After all, there are numerous types of covalent modifications such as methionine oxidation (Bender et al., 2008) that do not require immediate repair; only when a specific modification impairs catalysis, a compensatory response is warranted. The latter conditions would all be met by a redox signal related to superoxide.

In consequence, the observed transcriptional induction by low-dose MPP^+^ treatment in LUHMES cells was indeed surprisingly uniform as regards the different RCCs (Fig. 1). Complex III and IV genes were somewhat more induced than genes related to the actual site of inhibition, complex I (Fig. 1). Moreover, glycolytic genes were also selectively induced over a selection of other catabolic and anabolic gene groups (Fig. 8), apparently reflecting a purposeful, integrated transcriptional response. The TCA cycle, in contrast, was only marginally induced, which might be related to the fact that this pathway conducts epigenetic regulation through its own set of independent signals, namely succinate, fumarate, and 2-hydroxyglutarate (Arnold and Finley, 2023). The induction of the RCCs was reduced by approximately half after application of the radical scavenger PHT, which did not alter any of the MPP^+^-evoked metabolic changes affecting the pH, lactate, glucose, and NADH as well as the NAD^+^/NADH ratio (Fig. 2). These data support the concept of PHT as a non-pleiotropic, direct radical scavenger (Hajieva et al., 2009; Mocko et al., 2010) that was here employed at a selective 20 nM concentration, which is less than 1/10,000 of its typical toxic dose in cell culture (Moosmann et al., 2001). Notably, ATP levels were unaffected by the employed, 10 µM dose of MPP^+^, which is known to require significantly higher doses (EC_50(ATP)_ = 65 µM) to modify ATP levels under identical conditions as adopted in this study (Zhang et al., 2015). It is unclear whether a higher dose of PHT would have effectuated more than the observed ∼50% reversibility of induced RCC transcription. We would consider this to be very well possible, since radical scavengers acting towards short-lived species inevitably require high concentrations to achieve their maximum effect. On the other hand, the altered NAD^+^/NADH ratio may have plausibly contributed to the induction of part of the compensatory response through sensors like CtBP and, potentially, sirtuins (Anderson et al., 2017).

The chemical identity of the emanated redox signal is unknown at present. Several cues narrow down the set of possible species, though: (i) The signal is scavenged by rather low concentrations of PHT. In view of the established chemistry of PHT and its congeners (Moosmann et al., 2001; Farmer et al., 2017), this points at a radical species and excludes hydrogen peroxide as well as simple diamagnetic electrophiles like aldehydes. (ii) The signal should be closely related to complex I inhibition, which primarily yields superoxide radical anions (Brand, 2016) that are too inert to act as signal themselves (Winterbourn and Hampton, 2008). Still, the signal should be relatable to superoxide. (iii) The signal should be diffusible in the cytosol and potentially reactive enough to directly attack reactive cysteines on DNMT3B and SIRT1, both of which are known to possess such cysteines (Chen et al., 2012; Shao et al., 2014). Still, the signal must not be too reactive, which would limit its diffusibility and entail toxicity. This excludes, for instance, hydroxyl radicals. In summary, we propose that perhydroxyl radicals (i.e., protonated superoxide) would fulfill the named requirements best, as they are water soluble (Bielski and Cabelli, 1991), rapidly produced from superoxide in the more acidic cytosol (Saran and Bors, 1994), very reactive with chain-breaking antioxidants mechanistically related to PHT (Bielski and Cabelli, 1991), and of intermediate reactivity with cellular compounds like fatty acids and amino acids (Bielski and Shiue, 1978; Bielski et al., 1983). Concomitantly, they exhibit strong selectivity for cysteine when it comes to amino acids (Bielski and Shiue, 1978). Finally, their local production in the cytosol would steeply rise with dropping pH, e.g. in case of cytosolic lactate production (Saran and Bors, 1994), representing a physiologically desirable effect.

The deployment of an epigenetic mechanism to control the collective upregulation of functionally connected household genes that generally do not require much individual regulation appears expedient and reasonable (Deaton and Bird, 2011). The physically scattered genes encoding mitochondrially imported RCC subunits (Suppl. Fig. 2) are known to be co-regulated as a whole and on the level of the individual complexes, and they employ an overlapping set of rather few transcription factors (Lenka et al., 1998; van Waveren and Moraes, 2008). Moreover, they are GC-rich and contain CpG islands in approximately 80% of cases (van Waveren and Moraes, 2008). The specific mechanisms recruited by the here described redox signal encompass DNA methylation (Fig. 3) as well as histone acetylation (Fig. 4 and 6) and seem to be executed at least in part by DNMT3B and SIRT1. Both players have been linked to PD before: variants in the DNMT3B gene were (softly) associated with sporadic PD in populations from Brazil (Pezzi et al., 2017) and China (Chen et al., 2017). A loss of SIRT1 activity, but not total SIRT activity has been described for cortical tissue samples from patients with PD and other Lewy body diseases (Singh et al., 2017). PD-associated polymorphisms in the SIRT1 gene (Zhang et al., 2012; Maszlag-Török et al., 2021) have also been reported, along with reduced SIRT1 mRNA levels in peripheral blood cells (Maszlag-Török et al., 2021) and reduced SIRT1 protein levels in serum from PD patients (Zhu et al., 2021).

Notably, the use of an epigenetic signal with its inherently limited specificity may also result in potentially adverse off-target effects. For example, we have observed a significant epigenetic activation of LINE1 retrotransposons in LUHMES cells treated with different complex I inhibitors including MPP^+^ (Baeken et al., 2020). Moreover, the well-established epigenetic induction of SNCA transcription (Jowaed et al., 2010; Matsumoto et al., 2010; Yang et al., 2017) also recapitulated here (Suppl. Fig. 3) may likewise be classified as adverse and disease-promoting. Inspecting the response of other PD-associated genes (Day and Mullin, 2021) to complex I inhibition, a variety of potentially meaningful effects were observed, some of which were also redox-related. Beyond SNCA, especially PARK2, GBA, PARK7, and ATP13A2 were substantially modulated (Suppl. Fig. 3). Of these, the latter two modulations were significantly PHT-reversible, suggesting a redox-epigenetic mechanism of regulation as described herein. On the other hand, PARK2 and GBA were essentially neutral to PHT treatment despite being highly induced (PARK2) or repressed (GBA) by MPP^+^, illustrating the overlap of redox-dependent and redox-independent mechanisms after complex I inhibition. The strong repression of GBA and ATP13A2 following complex I inhibition as well as the induction of SNCA recapitulate the effects of hereditary, PD-causing mutations and duplications, respectively (Lesage and Brice, 2009). In particular, the known involvement of the GBA protein product (glucocerebrosidase) in α-synuclein degradation (Smith and Schapira, 2022) and the anti-aggregation properties of the ATP13A2 protein towards α-synuclein (Si et al., 2021) indicate that complex I inhibition exerts a triple, arguably synergistic effect towards toxic α-synuclein accumulation (Zharikov et al., 2015) through altered transcriptional regulation.

## Conclusion

Complex I-inhibited mitochondria emit an imminent demand signal to the nucleus in order to induce a broad-spectrum, but still statistically selective induction of mito-metabolic genes. The employed signal can be suppressed by a one-electron antioxidant which prevents the otherwise induced loss of DNA methylation, increase in histone acetylation and the reduction of SIRT1 and nuclear DNMT3B expression. The epigenetic regulation of retrotransposons and certain PD-related genes including SNCA following complex I inhibition may to constitute inadvertent side-effects of the physiologically purposeful, but arguably pleiotropic induction of mito-metabolic genes via epigenetic chromatin remodeling. The search for the elusive origin of complex I inhibition in PD should be intensified, as it appears to be upstream of the epigenetic alterations in the disease.

## Acknowledgements

The study was supported by the Corona foundation and the Manfred-und-Ursula-Müller foundation, members of the “Stifterverband für die Deutsche Wissenschaft”, DFG Grant CRC1177 and Volkswagen Foundation. The authors thank Dr. Ankush Borlepawar (Institute for Translational Medicine, Medical School Hamburg (MSH)) for critically reading the manuscript, and the team from QPS Austria for their careful planning and excellent performance of the animal experiments.

## Materials and Methods

### Chemicals and cell culture media

All cell culture media and supplements including Advanced Dulbecco’s Modified Eagle’s Medium (DMEM)/Ham’s F-12 (F12), Phosphate-Buffered Saline (PBS), N2 supplement, Hoechst 33258, and 6-diamidino-2-phenylindole (DAPI) were obtained from Invitrogen (Carlsbad, CA, USA). General laboratory chemicals and biochemicals, neurotoxins, phenothiazine and its derivatives were purchased from Sigma-Aldrich at the highest available purity unless otherwise specified.

### Cell culture

LUHMES cells were kindly provided by Dr. Jürgen Winkler (Division of Molecular Neurology, University of Erlangen, Germany). The cells were grown and differentiated as described (Krug et al., 2014; Baeken et al., 2020). At the 4^th^ day post differentiation, the medium was exchanged, and the cells were treated with different compounds at the following concentrations: MPP^+^: 10 µM; PHT: 20 nM; MPHT: 20 nM; APHT: 20 nM; 6-thioguanine: 1 µM; EX-257: 100 nM;

TSA: 50 nM.

### In vivo experiments

In vivo experiments in male C57Bl/6J mice (from Charles River) were performed by QPS Austria GmbH (Grambach, Austria). QPS Austria is accredited by the Association for Assessment and Accreditation of Laboratory Animal Care (AAALAC). Animal care, housing and experimentation were approved by the institutional Animal Care and Welfare Committee and complied to the animal welfare legislation of the Ministry of Science of the Austrian government.

Thirty mice aged 10 ± 2 weeks were allocated to three different treatment groups: a “vehicle group”, an “MPTP group”, and an “MPTP plus phenothiazine” group, involving repeated applications or injections of the two agents phenothiazine (10 mg/kg, ten doses in toto, vehicle DMSO:corn oil 1:50) and MPTP (20 mg/kg, four doses in toto, vehicle saline). The adopted treatment regimen is detailed in Figure 5 and has been described before (Baeken et al., 2020). Rotarod test: Animals from all groups were subjected to the Rotarod motor performance test on day 6 (one day after the final PHT treatment). Prior to the first test session, mice were habituated to the testing system until they were able to stay on the rotating rod at a constant speed of 2 rpm for approximately one minute. During testing, a single animal was exposed to the apparatus for three 180 s trials. The initial speed was increased from 2 rpm to 20 rpm during these 180 s. If the mouse fell, the session was over. The mean latency to fall in every single testing was determined.

Sample collection: After finishing the behavioral testing on day 6, the mice were deeply anesthetized by pentobarbital injection (600 mg/kg), transcardially perfused with saline, and dissected. The brains were hemisected, and the left hemisphere was subdivided into striatal tissue, midbrain and residual brain. These samples were rapidly frozen and kept at −80°C until further analysis. The right hemispheres were immersion fixed in freshly prepared 4% paraformaldehyde in PBS for 1 h at room temperature. Thereafter, they were transferred to 15% sucrose in PBS until they sank down, indicating sufficient cryoprotection. The hemispheres cryo-embedded in O.C.T. medium with dry ice-cooled isopentane and afterwards stored at −80°C.

### Immunohistochemistry

O.C.T. embedded brain hemispheres were cut into 10 µm slices with a cryostat. After washing with PBS, the sections were incubated with 3% BSA/0.1% Triton X-100 in PBS for 1 h at 4°C for blocking and permeabilization. After several washes with PBS, the slices were incubated overnight with primary antibodies at 4°C, repeatedly washed again, and secondary antibodies were applied for 2 h at RT. Following three more washing steps with PBS, 50 ng/ml Hoechst 33258 was added for 15 min at RT, before a final PBS wash. Finally, the sections were mounted in anti-fading solution (polyvinyl alcohol/p-phenylendiamine) and stored at −80°C until microscopic evaluation.

The following antibodies were employed: anti-tyrosine hydroxylase (ab112, Abcam, 1:1000 in PBS containing 1% BSA), anti-H3K14ac (7627, Cell Signaling, 1:100), and anti-SLC6A3 (DAT) (NBP2-22164, Novus Biologicals, 1:100). Secondary antibodies, Cy3-anti-rabbit and Cy2-anti-mouse (Jackson Immunoresearch) were used at 1:1000 dilution for tyrosine hydroxylase, and at 1:200 dilution for SLC6A3 and H3K14ac. All sections were visualized using standard fluorescence microscopy.

### Immunocytochemistry

LUHMES cells were plated at 7×10^4^ cells/cm² on glass cover slips with standard coating. The experiments were performed at day 6 post differentiation as described above. The procedure followed published protocols (Baeken et al., 2020; Baeken et al. 2021). Cells were incubated with primary antibodies overnight at 4°C and thereupon incubated with secondary antibodies (Cy3-anti-rabbit/Cy2-anti-mouse, 1:400) for 2 h at RT. Hoechst 33258 (50 ng/ml in PBS) was used as nuclear counterstain.

The following primary antibodies were used: anti-acetylated lysine (9441, Cell Signaling, 1:200), anti-DNMT1 (ab13537, Abcam, 1:100), anti-DNMT3A (2160, Cell Signaling, 1:100), anti-DNMT3B (67259, Cell Signaling, 1:100), anti-MAP2 (M4403, Sigma-Aldrich, 1:200), and anti-5-methylcytosine (28692, Cell Signaling, 1:200). Cy3-anti-rabbit/Cy2-anti-mouse (Jackson Immunoresearch), 1:400 were used as secondary antibodies. For the quantification, we followed published methodology (Baeken et al., 2020, Baeken et al. 2021). For DNMT1, DNMT3A and DNMT3B, nuclear localization was quantified through counting of positive nuclei normalized on total nuclei as stained with Hoechst 33258.

### Microscopy and image analysis

Slides generated through immunohistochemistry were recorded with an Axiovert 200 fluorescence microscope from Zeiss using blue, green and red filters and objectives for 4x and 10x magnifications. Immunocytochemistry slides were photographed with a laser scanning microscope (LSM) TCS SP5 from Leica (higher magnifications).

Immunohistochemistry pictures were evaluated using the open-source software ImageJ (imagej.net). Tyrosine hydroxylase (TH) staining was measured after the signal was „watershed“: this algorithm calculates signal maxima and thereby allocates cell borders. Cells were quantified using the “analyse particles” function from ImageJ that counts each continuous signal as one particle. Through this approach, cells can be evaluated regardless of the size of their cell body. For H3K14ac staining, total nuclei on the slide were quantified with DAPI, again using the “analyse particle” function of ImageJ. The same process was repeated for the H3K14ac staining, and the quotient H3K14ac/nuclei was calculated. Total acetyl-lysine and 5-methylcytosine levels were quantified by dividing the intensity measured with ImageJ by the number of cells (nuclei). Stainings of DNMT1, DNMT3A and DNMT3B were evaluated by counting the number of DNMT-positive nuclei and dividing it by the total number of nuclei indicated by DAPI.

### RNA sequencing and bioinformatics

To collect total RNA, LUHMES cells were harvested in 500 µl TRI-Reagent (Sigma-Aldrich) following the manufacturer’s instructions.

Residual DNA was removed by addition of 2 µl DNAse I (Thermo Fisher Scientific, #18047019) and incubation at 37°C for 1 h. Afterwards, 500 µl 75% ethanol were added, and the samples were washed again through centrifugation at 7,500g for 5 min at 4°C. The supernatant was discarded, and the cleaned RNA pellet was reconstituted in 50 µl DEPC-treated H_2_O.

The cleaned RNA was analyzed with a 2100 Bioanalyzer from Agilent and quantified using the Qubit dsDNA HS Assay Kit using a Qubit 2.0 Fluorometer (Life Technologies). All nine samples were pooled in equimolar ratio and sequenced in a NextSeq 500 High Output Flowcell with unique adapter sequences, SR for 1x 84 cycles, plus 7 cycles for the index read.

Sample demultiplexing and FastQ file generation was performed using Illumina bcl2fastq v2.19.1, and overall sequence quality was assessed with FastQC v.0.11.5. Sequence reads were aligned to the human reference genome GRCh38 with annotation from Gencode release 25 using STAR v.2.5.2b with parameters “--outFilterMismatchNmax 2 --outFilterMultimapNmax 10”. Secondary alignments were removed with SAMtools v.1.5, and data quality was examined using RSeQC v2.6.4 and dupRadar v.1.8.0. Read summarisation on the gene level was performed using Subread featureCounts v.1.5.1 with stranded option “-s 2”. Pairwise differential expression comparisons between sample groups were performed using the Bioconductor package DESeq2 v.1.18.1 using a cutoff of 1% FDR.

### Western blot analysis

Whole midbrain tissue and LUHMES cells were homogenized in lysis buffer (50 mM Tris-HCl, pH 6.8; 2% SDS; 10% sucrose; 0.5 mM EDTA, 0.5 mM EGTA plus protease and phosphatase inhibitor cocktails (Sigma-Aldrich)) and briefly sonicated. Protein concentrations were determined through BCA kit (Pierce) following the manufacturer’s protocol. A total of 20 µg protein were loaded on 12% SDS-PAGE gels and separated with a Mini protean III system (Bio-Rad). The proteins were transferred onto nitrocellulose membranes by electroblotting adopting standard protocols. Following 30 min incubation with 4% low-fat dry milk in PBST, the membranes were incubated with the following primary antibodies: anti-H3K14ac (7627, Cell Signaling, 1:1000), anti-acetylated lysine (9441, Cell Signaling, 1:1000), anti-DNMT3B (67259, Cell Signaling, 1:1000), anti-SIRT1 (8469, Cell Signaling, 1:1000), anti-SIRT3 (2627, Cell Signaling, 1:1000), or anti-SIRT4 (ab90485, abcam, 1:1000). Anti-α-tubulin (T9026, 1:1000, Sigma-Aldrich) and anti-histone H3 (14269, Cell Signaling, 1:1000) were used as controls for equal protein loading. Primary antibodies were generally detected with horseradish peroxidase-conjugated secondary antibodies (Jackson Immunoresearch). All antibodies used were diluted in PBST. Densitometric analysis of the immunoreactive bands was performed using the ImageJ software.

### Biochemical analyses

ATP, NAD^+^, NADH, glucose and lactate were measured using commercial test kits, following the instructions of the suppliers. Colorimetric or fluorimetric quantifications of the respective readouts were done in a multiwell plate reader (Wallac Victor 3 V, PerkinElmer, Waltham, MA, USA) and conducted in white 96-well plates.

For determination of cellular ATP levels, the medium was aspirated, and the cells were lysed in the supplied buffer. The employed ATP Detection Assay Kit (#700410 from Cayman Chemicals, Ann Arbor, MI, USA) uses a firefly luciferase system to translate ATP from cell and tissue lysates into luminescence.

NAD^+^ and NADH measurements were done with the NAD/NADH-Glo Assay Kit (#G9071 from Promega, Madison, WI, USA). Cells were grown in 24-well plates, washed, and lysed in 50 µl PBS/50 µl 0.2 M NaOH supplemented with 1% dodecyltrimethylammonium bromide (DTAB). A volume of 50 µl of the lysate was directly heated to 60°C for 15 min to eliminate NADH, the other 50 µl of the lysate were heated to 60°C for 15 min after adding 25 µl 0.4 M HCl to eliminate NAD^+^. To stabilize the pH, 25 µl of 0.5 M Tris, or 50 µl of 0.5 M Tris/HCl were subsequently added to the lysates. NAD^+^ and NADH levels were then quantified using the luciferin-coupled, NAD-cycling enzymatic test system provided with the kit.

Glucose and lactate in the medium were determined using the Glucose Colorimetric Assay Kit (#10009582 from Cayman Chemicals) and the Glycolysis Cell-Based Assay Kit (#600450 from Cayman Chemicals), respectively. Prior to analysis, pH values were documented with a calibrated mini-electrode device. To avoid interference of phenol red in the medium, a baseline background measurement was performed for each sample. For lactate measurements, 10 µl medium were used and diluted with water.

ROS levels were determined by fluorescence microscopy, coupled with image analysis. Cells grown on coverslips were treated as indicated and were incubated with 5 µM CellROX Deep Red (#C10422 from Invitrogen) for 30 min at 37°C under standard cell culture conditions. Following fixation with 4% PFA and counterstaining with 1 µg/ml DAPI, fluorescence microscopic images were taken from three separate fields of view with approximately 50 cells per image and analyzed with the ImageJ software.

### SIRT1 activity assay

SIRT1 activity in cells lysates was determined with the FLUOR DE LYS SIRT1 fluorometric drug discovery assay kit from Enzo Lifesciences (BML-AK555-0001). In brief, LUHMES cells were harvested in 200 µl SIRT1 assay buffer, briefly sonificated and estimated for protein content with a NanoDrop 1000 photometer. The highest possible amount of protein (181 µg) was diluted in SIRT1 assay buffer to a final volume of 35 µl for all samples. Four wells per sample were loaded with protein lysate and kept on ice for the remaining procedure. 64 µM of the SIRT1 substrate, FLUOR DE LYS SIRT1, and 500 µM NAD^+^ were diluted in 15 µl assay buffer and added to three of the four sample wells. The fourth sample received only 15 µl assay buffer to allow evaluation of the lysate background, while one well only received 50 µl SIRT1 assay buffer, and one well received sample buffer with NAD^+^ and FLUOR DE LYS SIRT1 to allow quantification of components’ background. The plate was incubated at 37°C for 1 h. The developer solution was prepared with SIRT1 assay buffer, 2 mM nicotinamide to stop additional reactions, and 1x FLUOR DE LYS Developer II. 50µl of the developer solution were added to each well, except for the background controls. Those wells instead received only 50 µl SIRT1 assay buffer. Two additional wells were prepared, one that only received 50 µl SIRT1 assay buffer and 50 µl developer solution, and one that received 64 µM FLUOR DE LYS Deacetylated Standard (in 50 µl assay buffer) and 50 µl developer solution, to allow quantification of the developer solution’s background as well as the potential signal maximum. The plate was then incubated at 37°C for 45 min. Afterwards, the fluorescent signal was quantified using a Multilabel Counter VICTOR 3V.

### Statistical analysis

All data are expressed as mean ± standard deviation (SD) of the indicated number of independent experiments. Statistically significant differences between the treatment groups were identified by either one-way or two-way ANOVA as indicated, followed by Benjamini-Hochberg multiple comparisons post-hoc test. Significance levels are either provided numerically or, in the case of RNA sequencing experiments, coded as symbols denoting p ≤ 0.05.

---

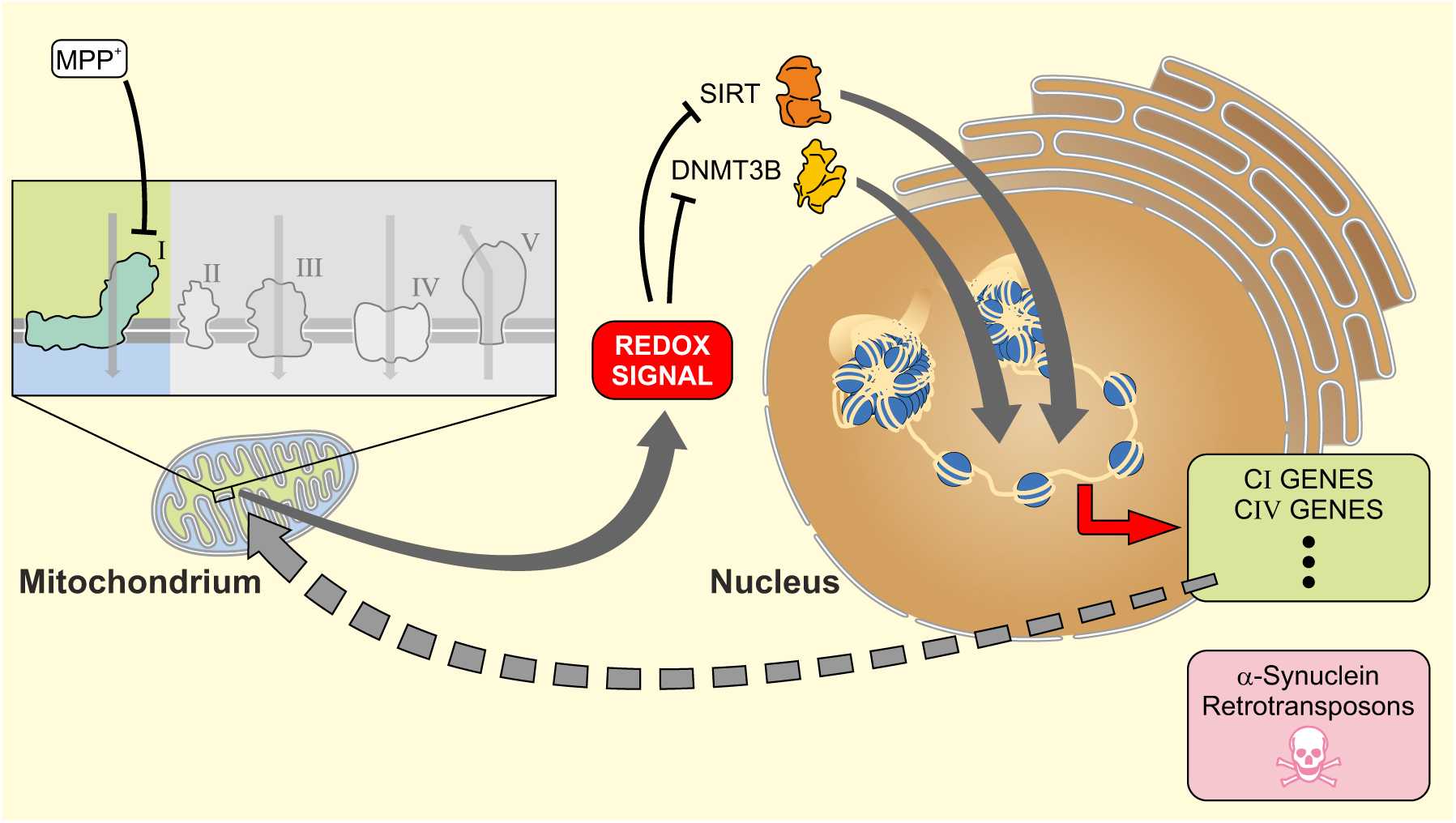

**Supplementary Figure 1.**
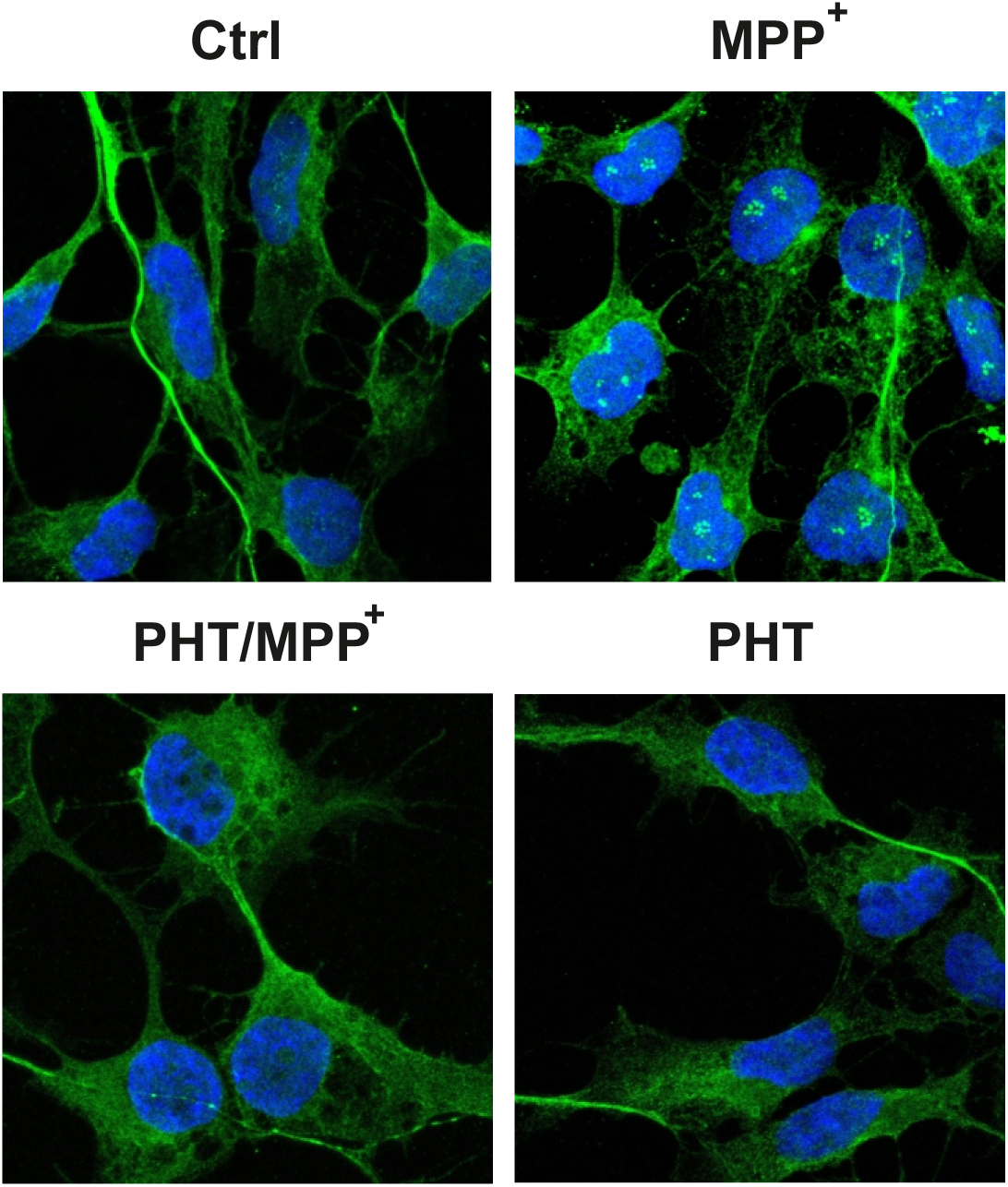
LUHMES morphology. Representative laser scanning microscopic images of differentiated LUHMES cells treated with 10 µM MPP^+^ or 20 nM PHT for 48 h (63x magnification). Microtubule-associated protein 2 (MAP2) is shown in green, DAPI in blue.

**Supplementary Figure 2.**
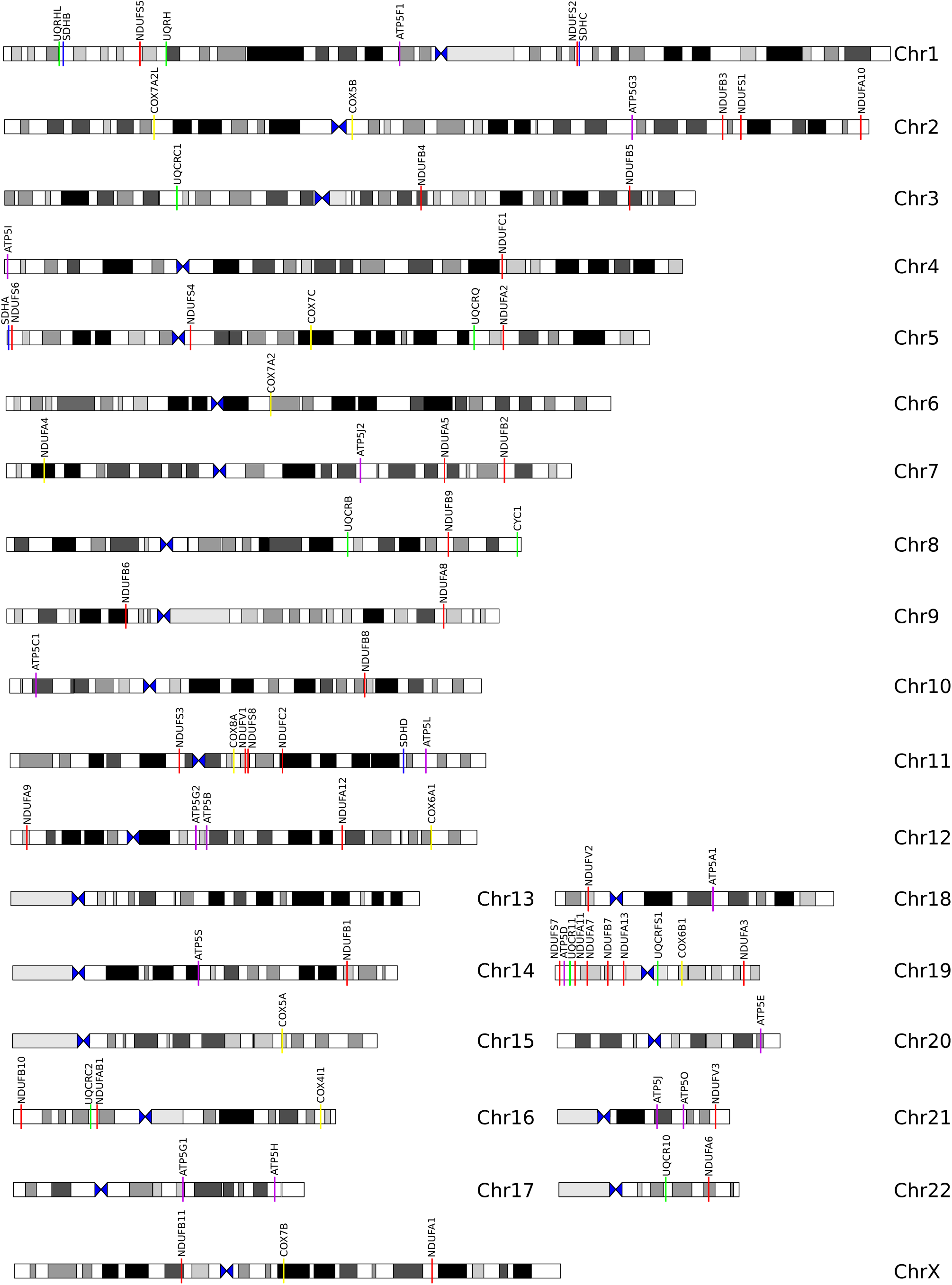
Genomic localization of genes coding for mitochondrially imported RCC subunits. The genomic site of origin of each of the analyzed RCC transcripts in Figure 1 is indicated using the same colour code (complex I red, complex II blue, complex III green, complex IV yellow, complex V violet).

**Supplementary Figure 3.**
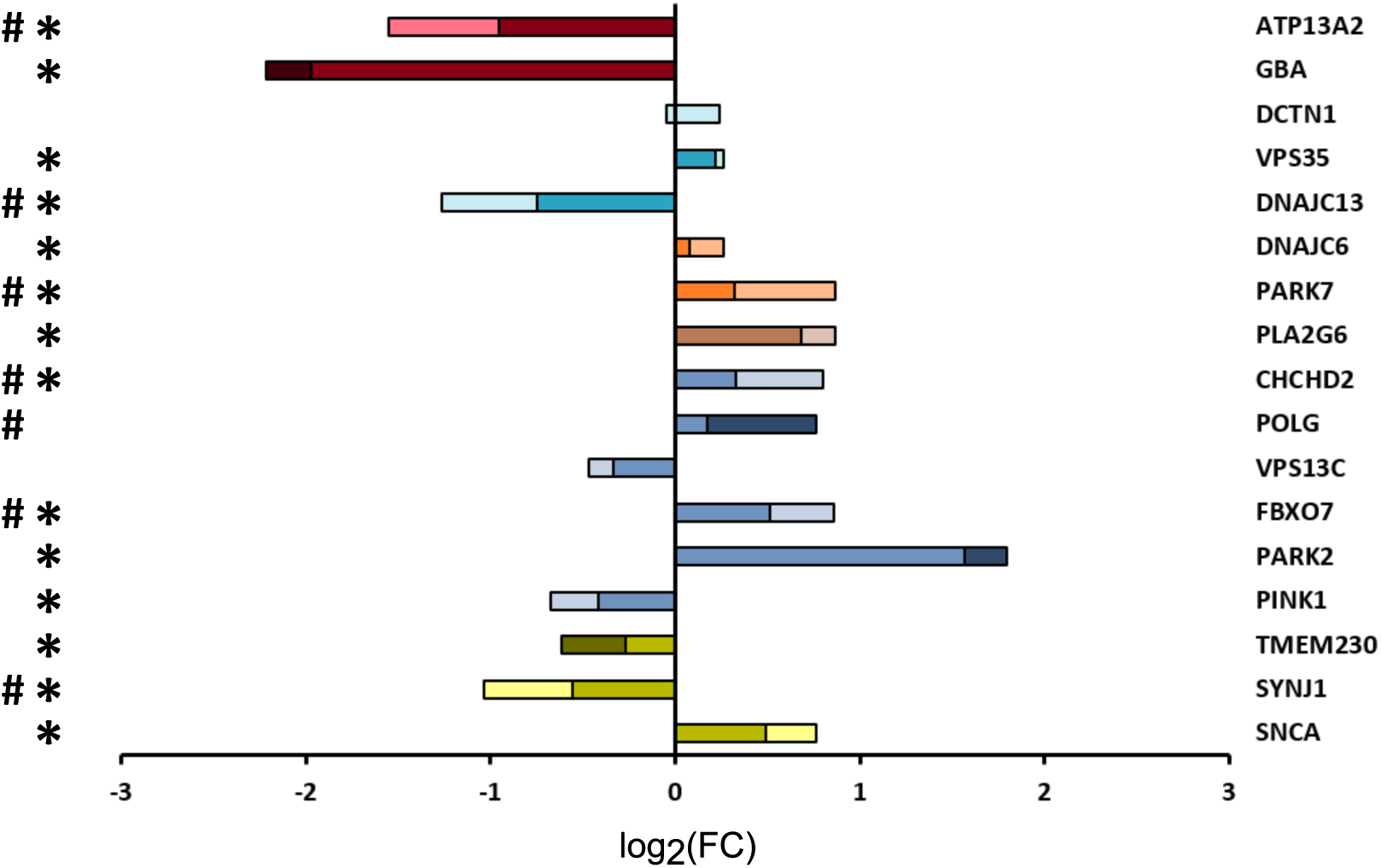
Transcriptional modulation of Parkinson’s disease-related genes by complex I inhibition in LUHMES cells. Regulation of PD-associated genes measured by RNA sequencing of differentiated LUHMES cells treated with 10 µM MPP^+^, 20 nM PHT or both for 48 h as in Figure 1. Transcriptional changes after MPP^+^ are indicated by bar length; regulation after PHT/MPP^+^ is indicated by color coding: lighter color denotes that the MPP^+^ effect was reduced by PHT, darker color denotes that the MPP^+^ effect was increased by PHT. Symbols indicate: *p-value ≤ 0.05 for MPP^+^ vs. control, ^#^p-value ≤ 0.05 for PHT/MPP^+^ vs. MPP^+^ by one-way ANOVA (n = 3). The genes were loosely classified by site and/or function, from top to bottom: lysosomal (red), retrograde transport (pale blue), chaperone (orange), fatty acid metabolism (brown), mitochondrial (blue), synaptic (green).

**Supplementary Table 1.**
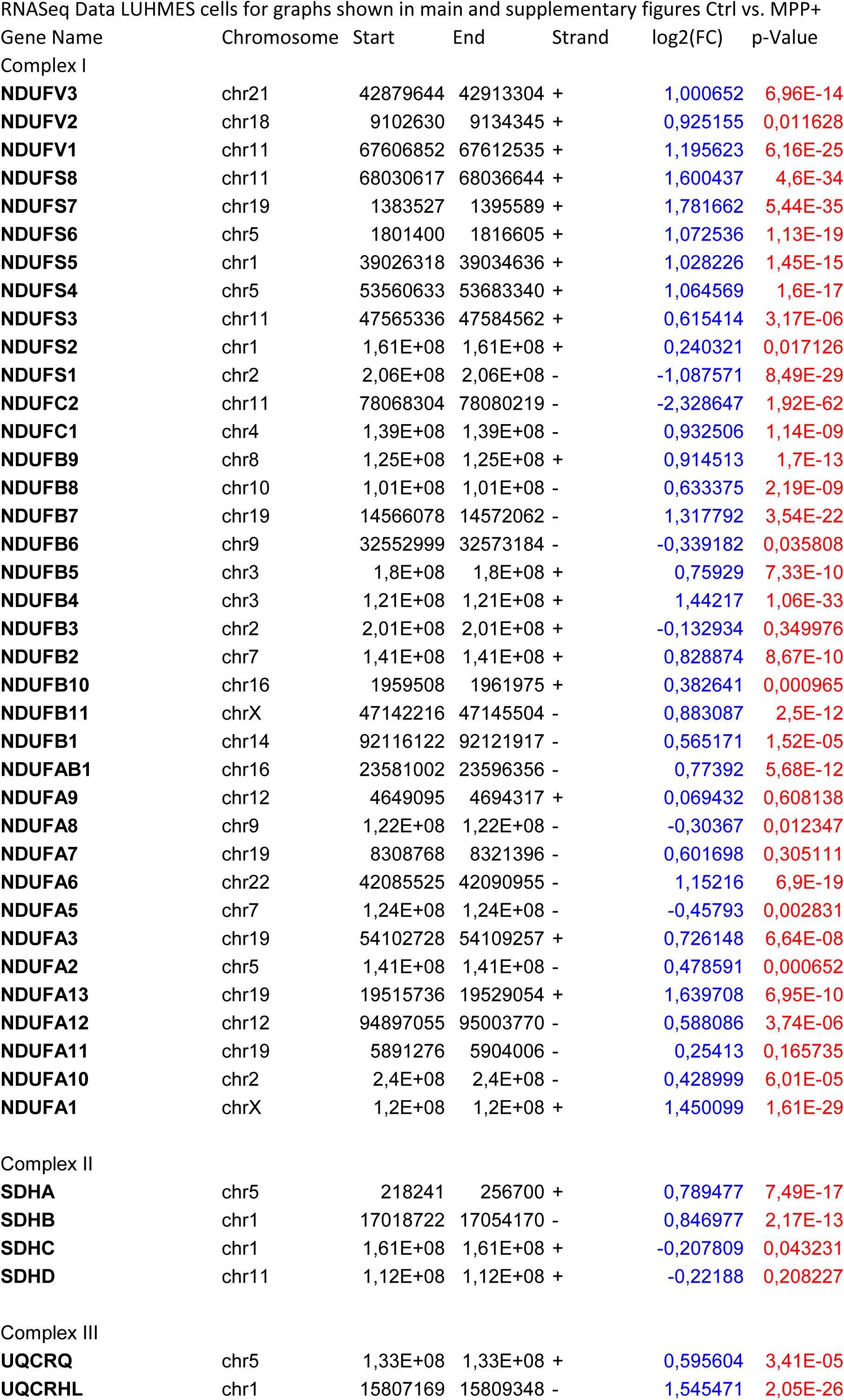
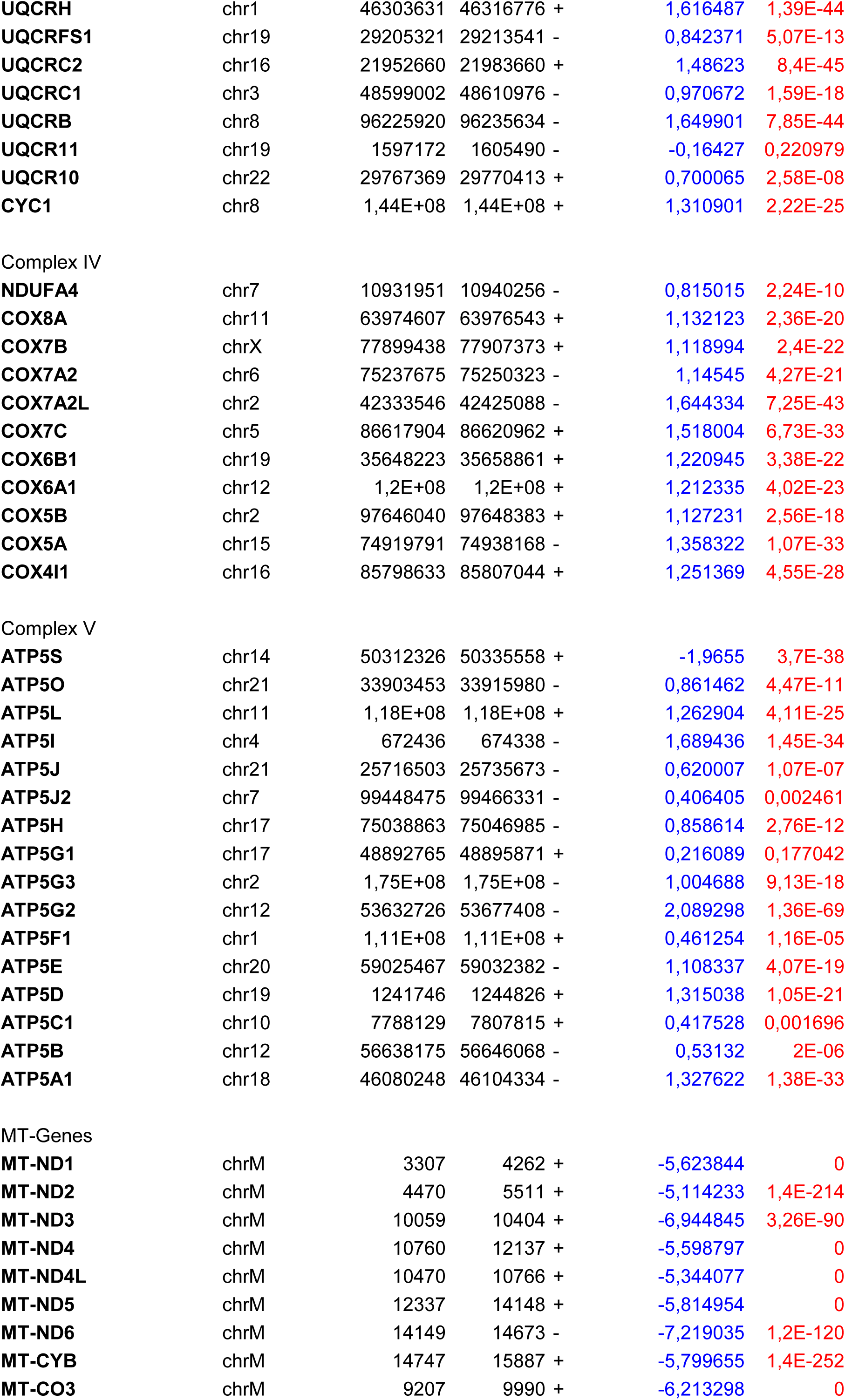
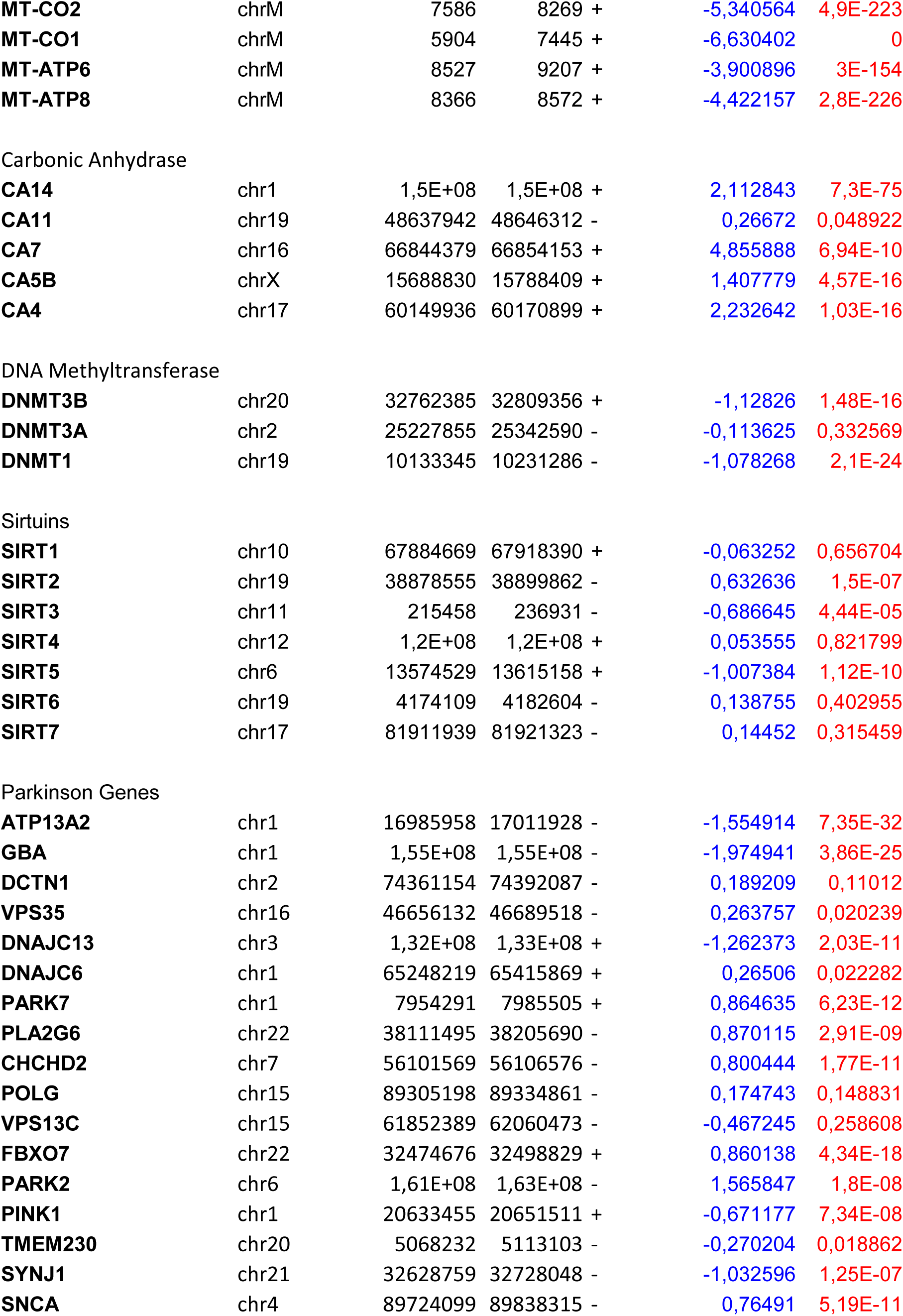
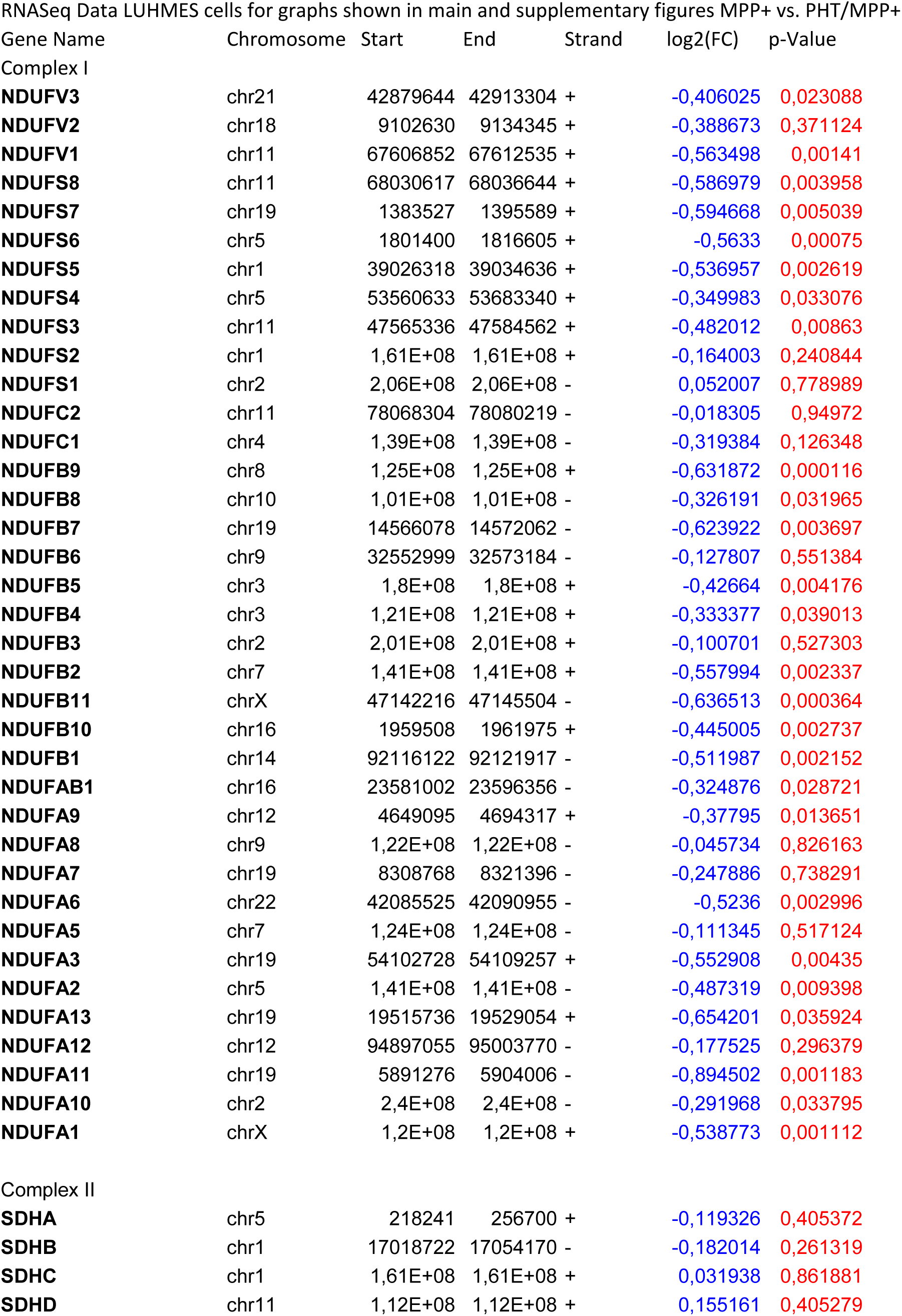
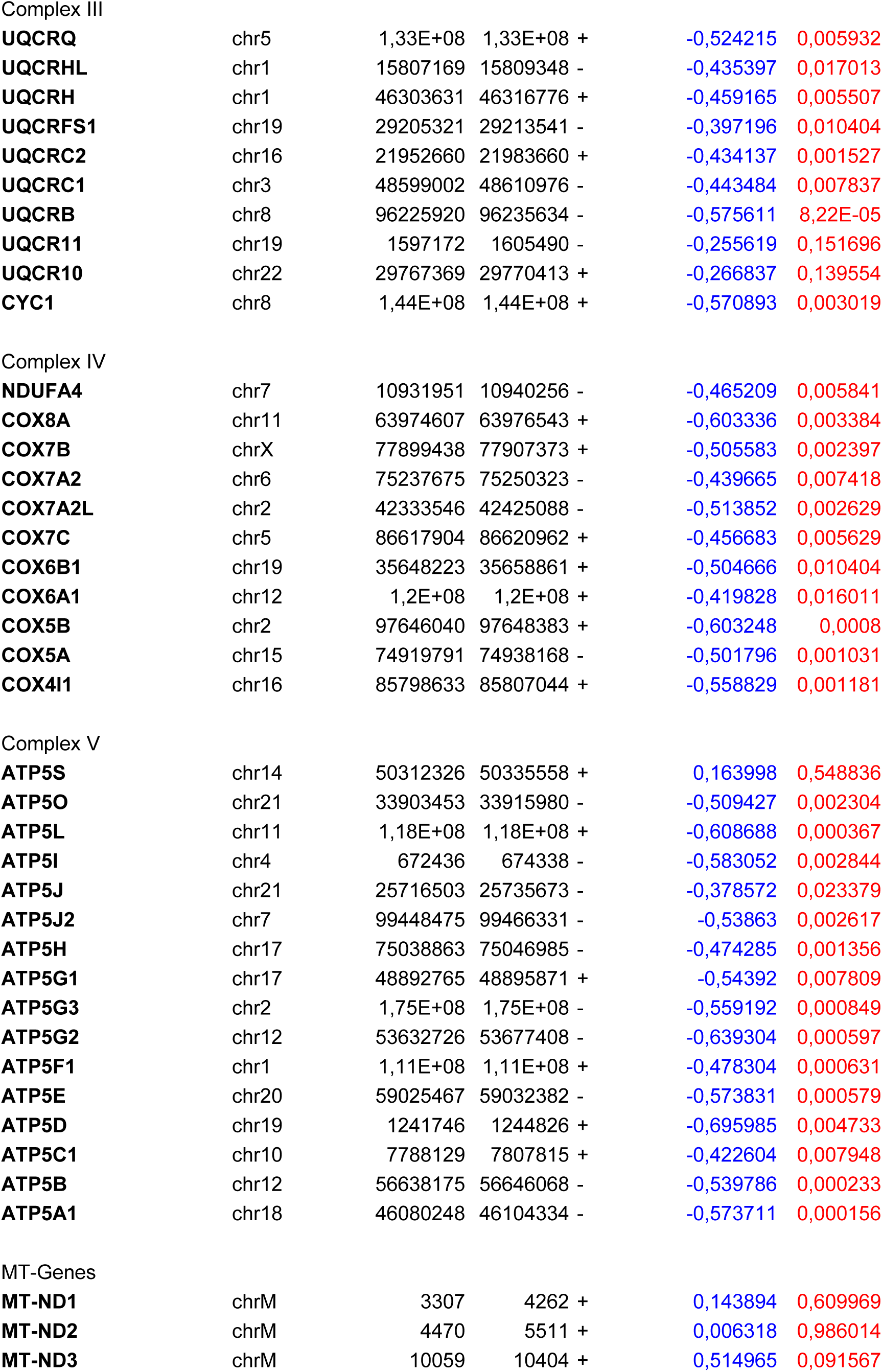
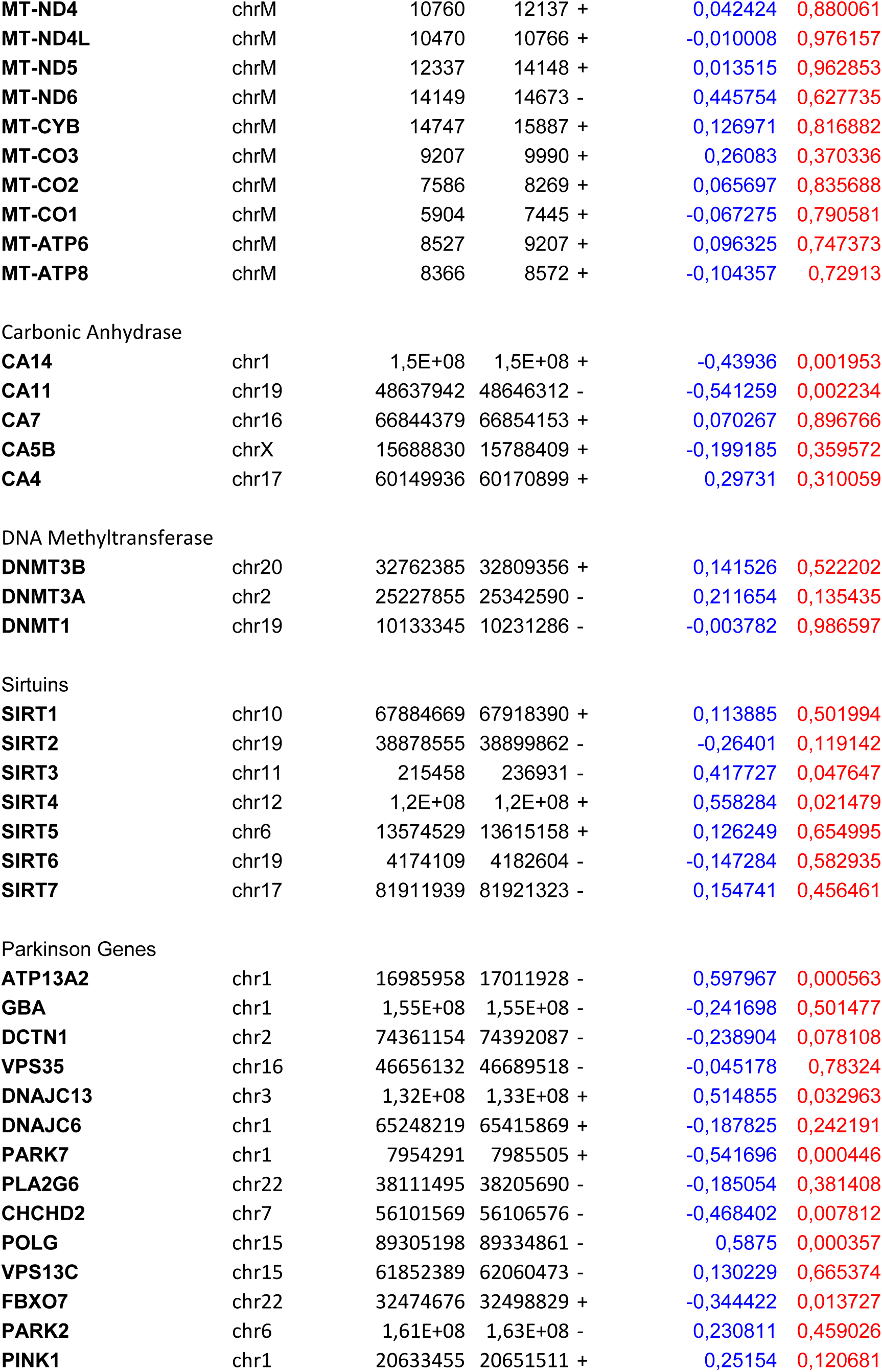
Chromosomal localization, fold change and statistical significance of the fold change of all genes displayed graphically in this work. The genes are listed in inverted alphabetical order and grouped to comprise complex I subunits, complex II subunits, complex III subunits, complex IV subunits, complex V subunits, mitochondrially encoded RCC subunits, carboanhydrases, DNA methyltransferases, sirtuins, and Parkinson’s disease related genes.

